# A thalamo-preoptic pathway promoting social touch

**DOI:** 10.1101/2022.01.11.475648

**Authors:** Dávid Keller, Tamás Láng, Melinda Cservenák, Gina Puska, János Barna, Veronika Csillag, Imre Farkas, Dóra Zelena, Fanni Dóra, Lara Barteczko, Ted B. Usdin, Miklós Palkovits, Mazahir T. Hasan, Valery Grinevich, Arpád Dobolyi

## Abstract

Social touch is an important form of communication, it is still unknown how it is processed. Here, we discovered a functional role for a neuronal pathway projecting from the posterior intralaminar thalamic nucleus (PIL) to the medial preoptic area (MPOA) in controlling social contact. Neurons in the PIL and the MPOA were activated by physical contact between female rodents and also by chemogenetic stimulation of PIL neurons. Chemogenetic stimulation of PIL neurons tagged by social contact experience increased direct physical interactions between familiar female rats without affecting other forms of social behavior. Furthermore, selective stimulation of the PIL-MPOA pathway, and the local activation of PIL terminals within the MPOA, elevated direct social contact between the animals suggesting the role of pathway-specific activated cell assemblies. Neurons projecting from the PIL to the MPOA contain the neuropeptide parathyroid hormone 2 (PTH2). The expression of the peptide was induced by social housing, the presence of PTH2 receptor was identified in MPOA neurons, and local injection of PTH2 increased the firing rate of identified preoptic area GABAergic neurons via the PTH2 receptor suggesting that PTH2 acts as a neurotransmitter in the PIL-MPOA pathway. We also found a homologous PIL to MPOA neuronal pathway in the human brain. Altogether, we discovered a direct thalamo-preoptic pathway, which bypasses the cerebral cortex and controls social touch. This pathway originates in neurons expressing PTH2, a neuropeptide recently shown in fish to respond to the social environment. These observations provide evidence for common evolutionary-conserved PTH2-containing social-touch specific engram circuits.

## Introduction

Social touch is a major component of social interactions. It is important in both sexual and nonsexual contexts in human (Ellingsen et al., 2015). In rodents, where modes of communication between conspecifics are more limited, the role of social touch may be even more pronounced, as part of the stereotypic behavioral repertoire for social interactions between individuals (Ebbesen and Froemke, 2021). The hypothalamus is a major regulatory center of rodent social behavior. It is also likely to be involved in the control of instinctive behaviors in humans (Zilkha et al., 2021). Social sensory inputs are known to reach the cerebral cortex via the thalamus, however, it is not known how information needed for social behavior arrives in the hypothalamus. Although, it is possible that information on social context reaches the hypothalamus via the cerebral cortex (Ahmadlou et al., 2021), it is also conceivable that the ascending sensory pathways carrying information on social touch might project directly to the hypothalamus. Within the responsible neuronal circuits, neuropeptides have been implicated in the control of a variety of social behaviors. Oxytocin, a pro-social neuropeptide, is known to promote social interactions, including social touch in rodents (Tang et al., 2020). Other neuropeptides, such as Tac2, were shown to play crucial roles in the behavioral response to chronic social isolation (Zelikowsky et al., 2018). Parathyroid hormone 2 (PTH2), a peptide originally named tuberoinfundibular peptide of 39 residues (TIP39) (Usdin et al., 1999), was recently demonstrated to sense the presence of conspecifics in zebrafish via mechanoreceptors in the lateral line organ (Anneser et al., 2020). In mammals, PTH2 expression is known to be elevated in mothers and participates in maternal care, as demonstrated by decreased motivation of mother rats to contact and suckle their pups after local delivery of a PTH2 receptor antagonist to the medial preoptic area (MPOA), located in the anterior hypothalamus (Cservenak et al., 2013). PTH2 is expressed in an unusual brain site, the thalamus, which otherwise is not rich in neuropeptides. Within the thalamus, PTH2 is confined to the posterior intralaminar thalamic nucleus (PIL), a multisensory information processing relay station (Dobolyi et al., 2018). Since PTH2 terminals are abundant in the MPOA, and PIL PTH2 neurons increased their activity in response to social interaction (Cservenak et al., 2017), we hypothesized that PTH2 neurons convey social inputs to the MPOA. To address this question, the expression of PTH2 in response to social interaction was first determined. Next, we examined the effect of chemogenetic-based stimulation of PIL neurons tagged by social contact experience. We found this to promote positively valenced physical contact between individuals (familiar female rats and mice were used to eliminate aggressive and sexual behaviors). Among the projection sites of the PIL, only the MPOA was exclusively activated by social touch and, in turn, selective stimulation of the PIL→MPOA pathway evoked direct physical contact between the animals. Thus, we have identified a new direct pathway, which controls social-touch driven social behavior independently from the established thalamo-cortical circuit.

## Results

### The PIL and its PTH2-exprsessing neurons

The PIL is located in the thalamus dorsal to the substantia nigra (SN) and ventromedial to the medial geniculate body (MG) (Fig. 1A). Borders of the PIL were defined using the location of PTH2 neurons as well as calcium-binding proteins. Parvalbumin immunoreactive (-ir) neurons were abundant in brain regions surrounding the PIL while only a few cells were labeled with parvalbumin within the PIL. In contrast, PTH2 and calbindin appeared exclusively in the PIL (Fig. 1B). The expression of PIL PTH2 mRNA was 2.4-times higher in animals kept in a social environment (together with conspecifics in their home cage) compared to control animals socially isolated for two weeks (Fig. 1C).

**Figure 1.**
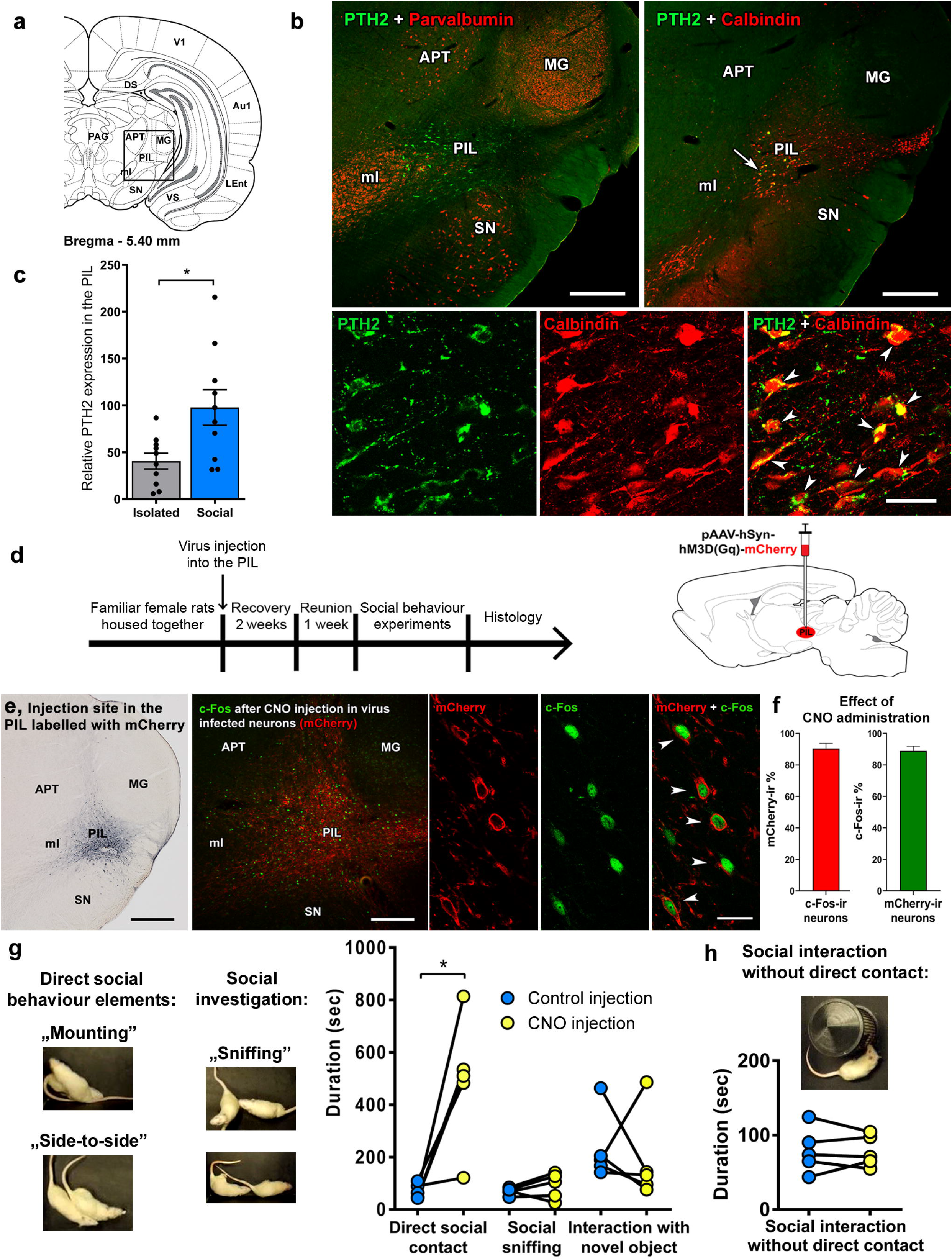
PTH2 content of PIL neurons and effect of chemogenetic stimulation of PIL on social behavior in rats. **a,** The PIL is located in the posterior part of thalamus in rat, dorsal from the SN and ventromedial from the MG as shown in a drawing of the rat brain (Paxinos and Watson, 2007). **b,** Chemoarchitecture of the PIL. Neurons in the PIL but not outside of it, highly expressed the neuropeptide PTH2 (green). There were only a few parvalbumin-positive neurons (red; upper left panel) in the PIL while parvalbumin-positive neurons were present in several surrounding brain areas, such as the anterior pretectal area (APT), the MG and the SN. In turn, calbindin-positive neurons (red; upper right and lower panels) were abundant in the PIL. Their distribution overlapped with that of the PTH2 neurons. High magnification confocal imaging demonstrated that PTH2 and calbindin are co-expressed in the PIL neurons (indicated by white arrowheads). Scale bars: 750 μm and 100 μm. **c,** Significantly higher PTH2 expression was present in the PIL in animals kept in a social environment in comparison with isolated controls, using quantitative RT-PCR. mRNA level of PTH2 / mRNA level of housekeeping gene * 10^6^, n=10 (each group), two-tailed unpaired t-test, \**p* = 0.013. Values are mean ± s.e.m. **d,** Experimental protocol of the chemogenetic stimulation of the PIL. **e,** The PIL as the injection site of the virus expressing mCherry fluorescent tag, and the *in vivo* validation of the effectiveness of chemogenetic stimulation. DREADD-expressing mCherry-immunopositive neurons (red) colocalized with c-Fos (green) in response to CNO injection. Scale bars: 750 μm (left), 300 μm (middle) and 60 μm (right). **f,** Quantification of CNO-induced c-Fos expression in the PIL. Most of the c-Fos expressing cells were mCherry-positive while most of the mCherry-positive neurons were activated after the injection of CNO, n=6. Values are mean ± s.e.m. **g,** The effect of chemogenetic activation of PIL neurons on the direct (physical) social contact between adult females during the half an hour recording. n=5, two-tailed paired t-tests, for direct social contact: *p* = 0.022, for social sniffing: *p* = 0.52, for interaction with novel object: *p* = 0.65; 0.010 < *\*p* < 0.050. **h,** The effect of the PIL stimulation on social interactions in the absence of physical contact (the animals were separated with bars). n=5, two-tailed paired t-test, *p* = 0.80.

### Stimulation of PIL elicits direct social contact in rats

#### The effect of chemogenetic activation of PIL neurons

To investigate whether PIL stimulation affects social behaviors, we injected pAAV-hSyn-hM3D(Gq)-mCherry into the PIL of female rats. The animals were subsequently housed individually for two weeks, and then reunited for 1 week (Fig. 1D) prior to injection of Clozapine-N-oxide (CNO), the agonist of the designer receptor exclusively activated by designer drug (DREADD). Almost all (90.4 ± 3.4%) activated cells contained mCherry and almost all (88.9 ± 3.0%) mCherry-positive neurons in the PIL showed c-Fos-positivity 1.5 hours after CNO administration, which validates the effectiveness of our chemogenetic stimulation (Fig. 1 E and F).

Next, we investigated several social behavioral elements following the stimulation of the PIL. The duration of direct social contact (such as mounting and side-to-side contact) significantly increased following stimulation of the PIL, compared with control injections (Fig. 1G). However, the duration of social investigation, estimated by the time of sniffing, and direct interaction with a novel object did not change. We also performed an experiment where direct contact between the animals was prevented by separating them by a bar wall which still allowed them to smell, hear and see each other (Fig. 1H). Chemogenetic activation of the PIL had no effect on the duration of social interaction under these conditions, highlighting the importance of direct contact as the effect of induced activation of the PIL.

#### Stimulation of PIL neurons, which were activated during previous social encounter

To selectively manipulate PIL neurons activated by social exposure, we used the **v**irus-delivered **G**enetic **A**ctivity-induced **T**agging of cell **E**nsembles (vGATE) method (Hasan et al., 2019). Excitatory and control viral cocktails (rAAV-(tetO)_7_-**P**_fos_-rtTA; rAAV-**P**_tet_bi-Cre/YC3.60; pAAV-**P**_hSYN_-DIO-hM3D(Gq)-mCherry/pAAV-**P**_hSYN_-DIO-mCherry) were injected into the PIL of female rats. In the vGATE system, a c-fos promoter (**P**_fos_) fragment drove the expression of the reverse tetracycline sensitive (tet) transactivator (rtTA). Following the recovery of the animals, a single intraperitoneal doxycycline administration launched the autoregulatory expression loop of (tetO)_7_-rtTA, thus opening the labeling period. The following day, the subjects were reunited with a cagemate in their home cage for 2 hours as direct social exposure, which resulted in the expression of the DREADD or control sequence in social contact experienced vGATE-tagged neurons only (encoded in the third virus: pAAV-**P**_hSYN_-DIO-hM3D(Gq)-mCherry or pAAV-**P**_hSYN_-DIO-mCherry) (Fig. 2A). Social behavior experiments were conducted 10 days later and then the brains were processed for histological analysis. Animals in diestrus were used for the behavioral experiments.

**Figure 2.**
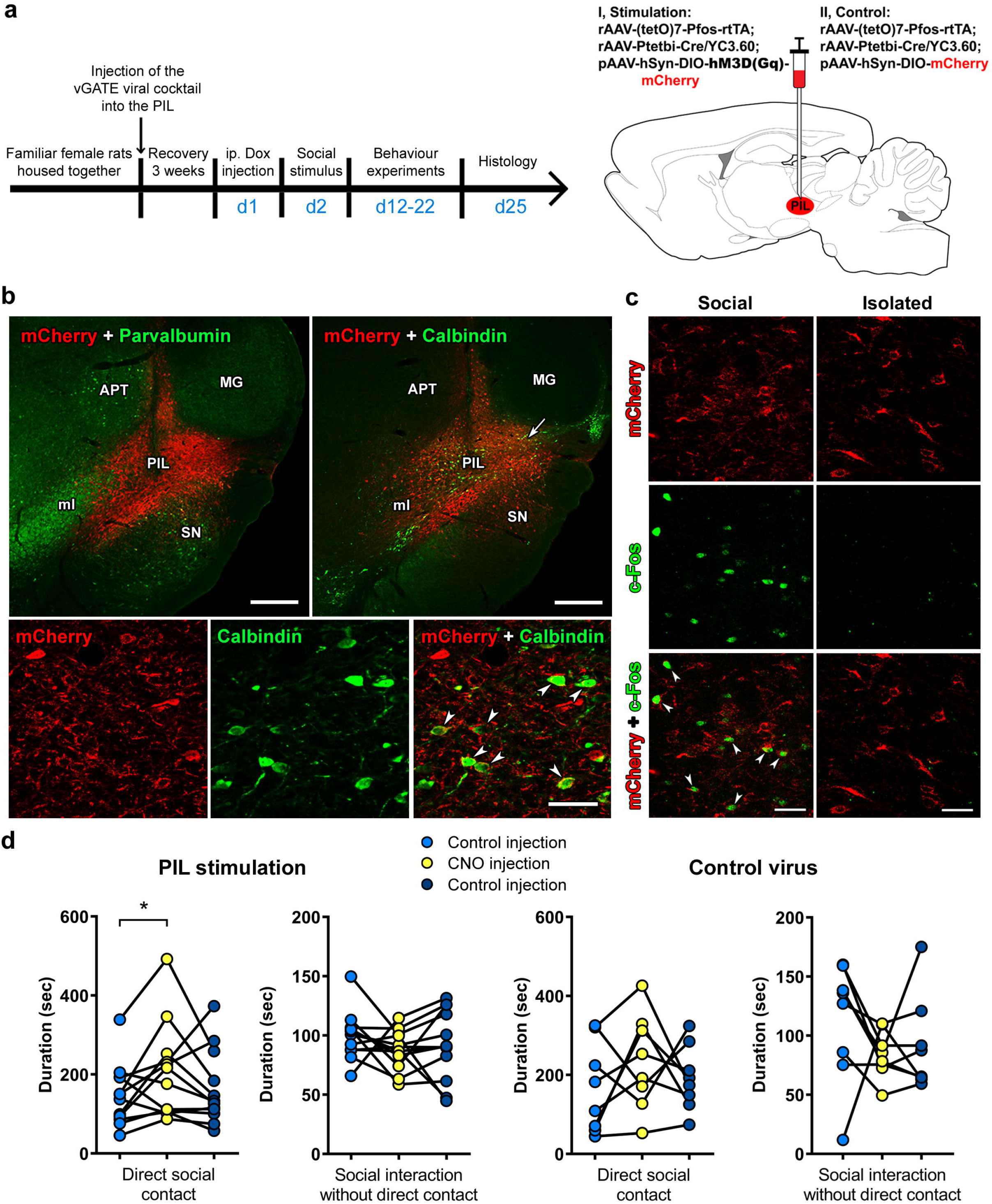
Effects of the stimulation of PIL using vGATE viral cocktail. **a,** Experimental protocol of the chemogenetic PIL stimulation using vGATE viruses. **b,** Distribution of PIL neurons infected by the vGATE viral cocktail. Parvalbumin and calbindin (green) indicate the borders of the PIL. High magnification confocal imaging demonstrates that calbindin was co-expressed with the mCherry fluorescent tag (red) in several neurons (indicated by white arrowheads) in the PIL. Scale bars: 500 µm (upper) and 100 µm (lower). **c,** *In vivo* validation of the vGATE technique: neurons infected by the vGATE viruses were activated after social exposure (indicated by white arrowheads) but not in isolated control animals. Scale bars: 75 µm. **d,** Social behavior experiments using the vGATE viral cocktail. The animals were placed in the same cage where they could freely interact with each other or were separated with bars preventing their physical contact. For PIL stimulation: n=12, two-tailed paired t-tests, for direct social contact: first control injection *vs* CNO: *p* = 0.041, CNO *vs* second control injection: *p* = 0.18; for social interaction without direct contact: first control injection *vs* CNO: *p* = 0.15, CNO *vs* second control injection: *p* = 0.59. For control virus: n=8, two-tailed paired t-tests, for direct social contact: first control injection *vs* CNO: *p* = 0.25, CNO *vs* second control injection: *p* = 0.47; for social interaction without direct contact: first control injection *vs* CNO: *p* = 0.22, CNO *vs* second control injection: *p* = 0.63; 0.010 < *\*p* < 0.050.

Labeling with calcium-binding proteins revealed that the PIL region infected with the vGATE viruses was surrounded with areas with parvalbumin-positive cells, and that the vast majority of the infected cells were calbindin-positive as PTH2-positive cells are (Fig. 2B). To validate the applied vGATE method, we performed c-Fos immunolabeling after direct social exposure or isolation. 2.9-times more calbindin cells were labeled for c-Fos after social exposure than after isolation, and 70.5% of the c-Fos-ir cell were mCherry-positive (representing vGATE labeled neurons) compared with 46.3% in the isolated group. In addition, 53.3% of the mCherry cells were c-Fos-positive after social interaction compared with 21.4% after isolation (Fig. 2C). During the direct contact social behavior test, where the animals could freely interact with each other, the duration of direct social contact significantly increased following the stimulation of vGATE-tagged neurons of the PIL, but no behavioral changes were found in the group injected with the control virus (Fig. 2D). Notably, only direct physical contact elicited an increase in social interactions time after chemogenetic activation of vGATE-tagged PIL neurons.

### Projections of the PIL and the activation of its target area during direct contact social behavior in rats

To determine neuronal connections of PIL neurons, the anterograde tracer biotinylated dextran amine (BDA) was injected into the PIL of female rats (Fig. 3 A and B). Labeled axons were found in a number of socially-relevant brain regions, such as the infralimbic cortex (ILC), lateral septum (LS), medial preoptic nucleus (MPN), medial preoptic area (MPA), ventral bed nucleus of stria terminalis (vBNST), paraventricular hypothalamic nucleus (PVN), amygdala (A), dorsomedial hypothalamic nucleus (DMH), periaqueductal central grey (PAG), and lateral parabrachial nucleus (LPB) (Fig. 3 C and D). Semi-quantitative analysis of labeled fibers revealed that the projections from the PIL have a distribution similar to that of a subset of PTH2-positive fibers, and that within regions the receive PIL projections the pattern closely resembles that of PTH2-positive fibers (Fig. 3E; SI Appendix, Table S1). A further aim was to find out whether social interaction activates the target areas of the PIL together with the activation of the PIL itself, which was reported previously (Cservenak et al., 2017). We found four brain regions, the ILC, MPOA, MeA and DMH, where the numbers of activated neurons were significantly higher after reuniting the rats after 1 week of social isolation as compared to isolated animals (SI Appendix, Fig. S1).

**Figure 3.**
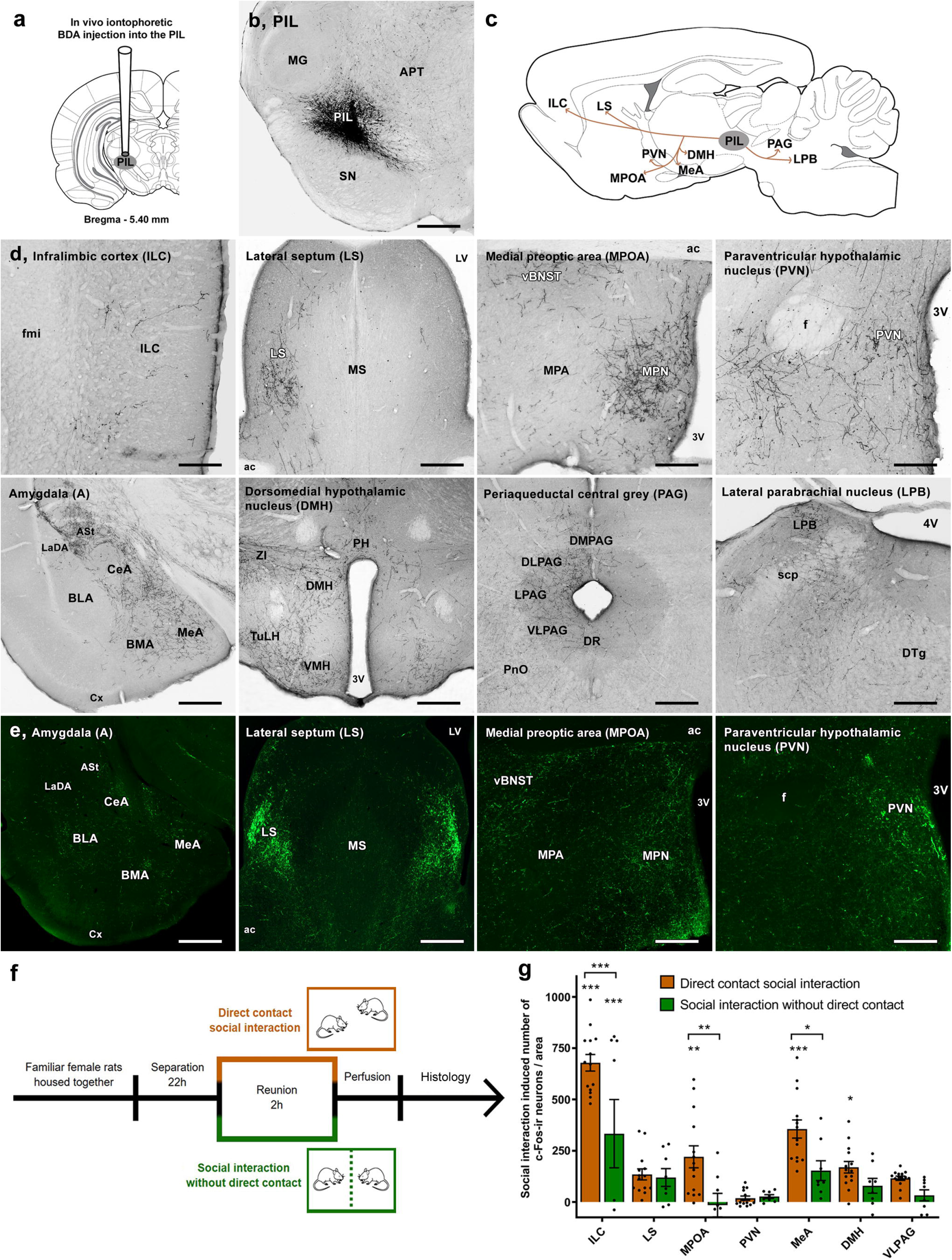
Anterograde projections of the PIL and its PTH2-expressing cells, and activation of PIL target area neurons upon social interaction with or without direct physical contact between the female rats. **a,** Injection of the anterograde tracer BDA into the PIL. **b,** A representative BDA injection site in the PIL. Scale bar: 750 μm. **c,** Summary of the projections of PIL neurons based on anterograde tract tracing. **d,** Anterogradely labelled fibres are shown in several brain regions implicated in social behaviour, such as the ILC, LS, MPOA, PVN, amygdala, dorsomedial and some other hypothalamic nuclei, PAG and some pontine structures including the LPB. Scale bars: 300 μm (ILC, PVN), 375 μm (MPOA), 450 μm (LPB), 600 μm (LS), 750 μm (A, DMH, PAG). **e,** PTH2-positive fibres in PIL target areas. The distribution of the PTH2-cointaining fibres is similar to the projections of the PIL as seen in the anterograde tracing. **f,** The experimental protocol for studying neuronal activation in response to social interaction performed in pair housed female rats. Rats were isolated for 22 hours before the social experiment to reduce basal c-Fos activation. The animals were reunited either in the same cage where they could freely interact with each other, or separated with bars preventing direct physical contact. The control animals remained isolated until perfusion. At the end of the 2 hour reunion (or final isolation period for controls), the rats brains were processed for c-Fos immunohistochemistry. **g,** Comparison of the c-Fos activation of seven socially implicated brain regions. The diagram represents the numbers of social interaction induced c-Fos-ir neurons (number of activated neurons after social exposure – number of activated neurons in isolated animals) / area. For direct contact social interaction: n=14, for social interaction without direct contact and isolated control group: n=8, two-way ANOVA followed by Šídák’s multiple comparisons tests. Direct contact social interaction *vs* social interaction without direct contact: for ILC: *p* < 0.001, for LS: *p* > 0.99, for MPOA: *p* = 0.0080, for PVN: *p* > 0.99, for MeA: *p* = 0.034, for DMH: *p* = 0.80, for VLPAG: *p* = 0.86; 0.010 < \**p* < 0.050, 0.001 < \*\**p* < 0.010 and \*\*\**p* < 0.001. Values are mean ± s.e.m.

We also examined the anterograde transport of the pAAV-hSyn-hM3D(Gq)-mCherry virus which was used to stimulate the PIL. mCherry-positive axons had the same distributional pattern as that found by traditional anterograde tracing (SI Appendix, Fig. S2A), which verifies the accuracy of both the injection sites and the anterograde transport of the virus. Based on the density of the labeled axons the MPOA and MeA appeared to be the main target areas of the PIL. Following control injections of the virus into the adjacent substantia nigra the labeled fibers were present only in the caudate-putamen, and not in the target areas of the PIL (SI Appendix, Fig. S2B).

In the chemogenetic experiment, the administration of CNO without social stimuli resulted in c-Fos activation in the PIL, and also in socially relevant target areas of PIL neurons in a pattern similar to that detected after social interaction (SI Appendix, Fig. S3A). Significantly higher number of c-Fos-positive neurons was found in the PIL, due to the direct effect of the chemogenetic stimulation, and in the PIL targets ILC, MPOA and MeA, compared to vehicle injection (SI Appendix, Fig. S3B).

The effect of direct physical contact was also determined in PIL target areas by comparing freely interacting rats with ones separated by bars (Fig. 3F). Three brain regions were identified in which the number of activated neurons was significantly lower when physical contact was prevented: the ILC, MPOA and MeA (Fig. 3G). With the exclusion of physical contact, the activation of the MPOA was similar to the control animals, without any social stimulus, highlighting the importance of direct contact in the activation of this particular region. Notably, one of the highest densities of PIL neuron axonal terminals was found in the MPOA (SI Appendix, Fig. S2A), so we further investigated the role of the PIL→MPOA circuit in the regulation of physical contact during social behavior using two different approaches.

### Projections of PIL neurons to the MPOA facilitates the behavioral effect of physical contact

A retrogradely transported virus expressing Cre-recombinase and mCherry fluorescent protein was injected into the MPOA. Three weeks later, a Cre-dependent virus expressing mCitrine and an excitatory DREADD was injected into the PIL. As demonstrated by expression of mCherry and mCitrine, respectively, the MPOA injected Cre-recombinase expressing virus spread to afferent brain areas including the PIL (Fig. 4 A and B), and the Cre-dependent mCitrine fluorescent protein and associated excitatory DREADD were expressed only in MPOA projecting PIL neurons (Fig. 4C). During a social behavior experiment, the selective chemogenetic activation of MPOA projecting PIL neurons increased the duration of direct social interaction (Fig. 4D). Following vehicle-only injections, a few days later, the duration of social behavior remained at basal level.

**Figure 4.**
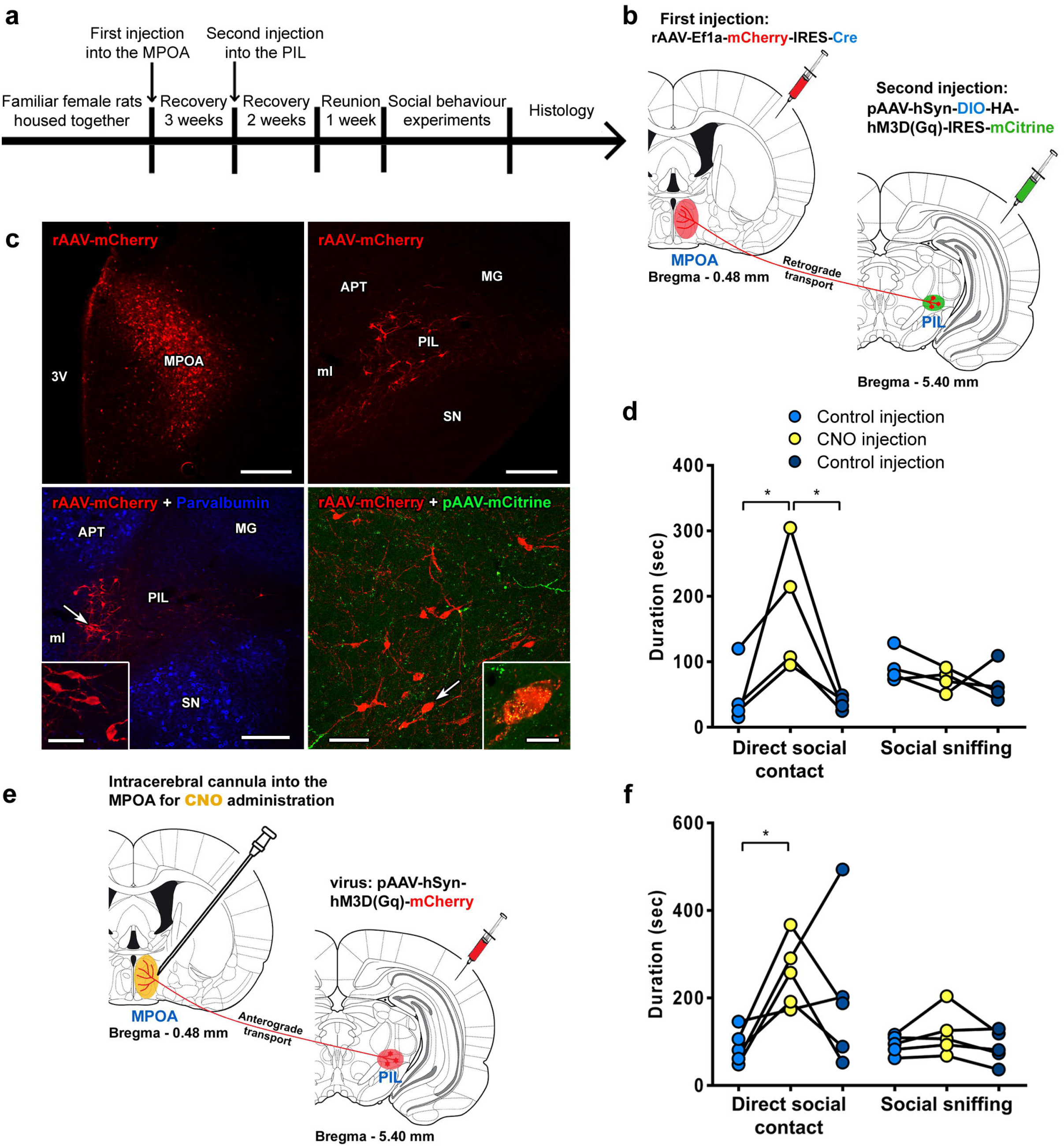
Demonstration of the role of PIL-MPOA projection in social touch. **a,** Experimental timeline for the selective chemogenetic stimulation of PIL neurons projecting to the MPOA. **b,** Experimental design of the double viral injections. **c,** The first two images demonstrate the injection site in the MPOA and the retrogradely labelled neurons in the PIL. The second two images show that the retrogradely labelled cells (red) are located in the medial PIL surrounded by parvalbumin-ir neurons (blue) and express the Cre-dependent DREADD (green). Scale bars: 400 μm (first two images), 300 μm and 100 μm (third image and its inset), 150 μm and 30 μm (fourth image and its inset). **d,** Effect of the selective chemogenetic stimulation of neurons projecting from the PIL to the MPOA on social behaviour. The animals were placed in the same cage where they could freely interact with each other. n=4, two-tailed paired t-tests, for direct social contact: first control injection *vs* CNO: *p* = 0.037, CNO *vs* second control injection: *p* = 0.028; for social sniffing: first control injection *vs* CNO: *p* = 0.059, CNO *vs* second control injection: *p* = 0.42; 0.010 < *\*p* < 0.050. **e,** Experimental design of the stimulation of fibre terminals of PIL neurons in the MPOA with local CNO administration using intracerebral cannula implanted into the MPOA. **f,** The effect of the selective chemogenetic stimulation of fibre terminals of PIL neurons in the MPOA on social behaviour. The animals were placed in the same cage where they could freely interact with each other. n=5, two-tailed paired t-tests, for direct social contact: first control injection *vs* CNO: *p* = 0.018, CNO *vs* second control injection: *p* = 0.27; for social sniffing: first control injection *vs* CNO: *p* = 0.20, CNO *vs* second control injection: *p* = 0.12; 0.010 < *\*p* < 0.050.

Chemogenetic stimulation of PIL axonal terminals in the MPOA was also performed using local delivery of CNO into the MPOA (Fig. 4E). The chemogenetic activation of axonal terminals of PIL neurons in the MPOA increased the duration of direct social interaction of the animals (Fig. 4F).

No significant effects were observed in other, non-contact, behavior tests performed following the stimulation of the PIL, including a social novelty preference test (SI Appendix, Fig. S4) and depression- and anxiety-like behavioral tests (SI Appendix, Fig. S5), implying that the function of the PIL is specific for the regulation of direct contact social behavior. We also found no social behavioral changes after the inhibition of the PIL (SI Appendix, Fig. S6) indicating that PIL is sufficient but not necessary for social contact behavior.

### Characteristics of PTH2-sensitive preoptic neurons

The parathyroid hormone 2 receptor (PTH2R) is the receptor for PTH2. Using PTH2R-ZsGreen mice for visualization of the receptor’s distribution, we found it to be abundant in the MPOA. Furthermore, the number of MPOA c-Fos positive PTH2R-expressing neurons was significantly higher after social interaction than in isolated control animals (Fig. 5A).

**Figure 5.**
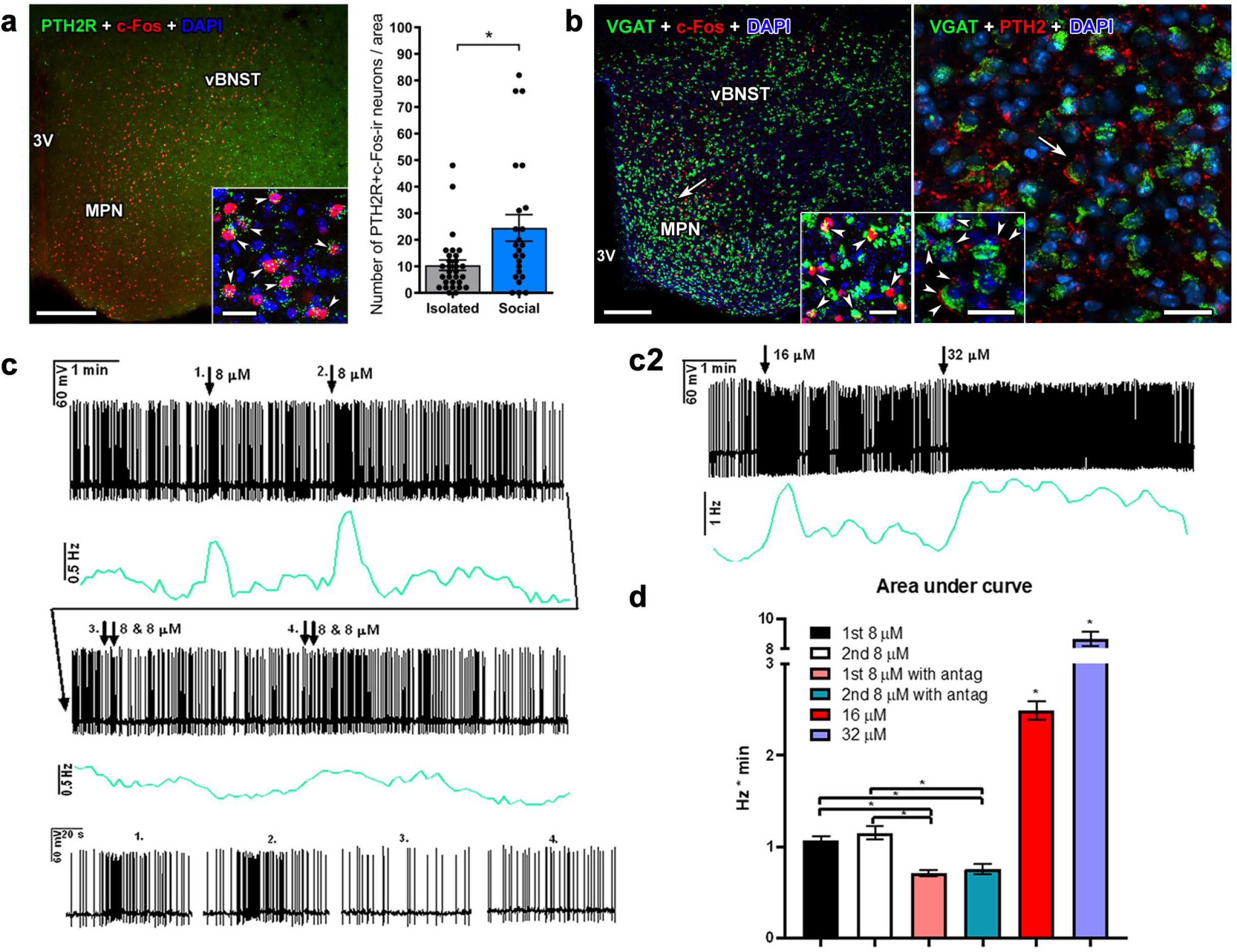
The effect of PTH2 on preoptic neurons. **a,** The PTH2R (green), the receptor for PTH2, was expressed in c-Fos activated (red) neurons (blue) of the MPOA after social contact in PTH2R-ZsGreen mice. The number of PTH2R/c-Fos positive neurons in the MOPA was significantly higher after social interaction with another female conspecific compared with isolated controls. n=30 for isolated, n=24 for social, two-tailed unpaired t-test, \**p* = 0.013. Values are mean ± s.e.m. Scale bars: 300 μm (main) and 50 μm (inset). **b,** Preoptic GABAergic (green) neurons expressed c-Fos following social exposure of VGAT-ZsGreen mice (left panel). PTH2-ir fibres (red) closely apposed the cell bodies (blue) of VGAT-positive neurons in the MPOA (right panel). Scale bars: 400 μm and 50 μm (left and its inset), 70 μm and 20 μm (right and its inset). **c1,** The neuronal activity of MPOA GABAergic neurons in female mice was enhanced by PTH2. The figure shows representative recordings during repetitive PTH2 applications that resulted in transient (less than 1 min) elevations in firing rate. The greenish lines under the recordings represent frequency distribution, presenting two peaks following the applications of PTH2. The recording of the same neuron continued further on (long arrow line). Here, double arrows mark the local application of a PTH2R antagonist immediately before PTH2 (both applied in 8 µM concentration) demonstrating that in the presence of the PTH2R antagonist PTH2 did not have an effect on the activity of the recorded neuron. The four insets in the bottom of the panel are 1 min magnified periods of the recordings, corresponding to the four applications of PTH2 in the absence (1^st^, 2^nd^) or in the presence (3^rd^, 4^th^) of the PTH2R antagonist. **c2,** Higher doses of PTH2 resulted in higher and longer responses, demonstrating dose dependency of PTH2 action on the firing rate of preoptic GABAergic neurons. **d,** Quantitative analysis of PTH2 action on the firing rate of preoptic GABAergic neurons. The effect of local bath application on neuronal activity was calculated as the area under the curve in the frequency distribution. Due to the different durations of the transient effect of the various doses of PTH2, the first four bars (with 8 µM) were calculated with 1 min time window, the 5th bar (16 µM) was calculated with 1.5 min window, the 6th bar (32 µM) was calculated with 5 min window. Area under curve: 1^st^ 8 μM: 1.17 ± 0.017 Hz*min, n=9; 2^nd^ 8 μM: 1.24 ± 0.014 Hz*min, n=9; 1^st^ 8 μM with antagonist: 0.58 ± 0.012 Hz*min, n=9; 2^nd^ 8 μM with antagonist: 0.61 ± 0.011 Hz*min, n=9; 16 μM: 2.49 ± 0.103 Hz*min, n=9; 32 μM: 8.67 ± 0.501 Hz*min, n=9. Repeated measures ANOVA was used to examine whether the firing rate change of each group was significant, * *p* < 0.001. Values are mean ± s.e.m.

We further investigated the connection between the GABAergic neurons in the MPOA and the PTH2 neuropeptide. We confirmed that GABAergic cells in the MPOA were activated after social exposure. Moreover, these cells were closely apposed by PTH2-positive fibers (Fig. 5B). Finally, we demonstrated with the patch-clamp technique used in acute slices that GABAergic

MPOA neurons of female VGAT-ZsGreen mice were activated by PTH2, as PTH2 applications resulted in dose dependent transient elevations in the firing rate of preoptic GABAergic neurons while an antagonist of the PTH2R eliminated this action (Fig. 5 C and D).

### The PIL-MPOA pathway in human brain

To investigate whether the PIL→MPOA circuit observed in rodents also exists in human brain we used autopsy material for histological analysis. Using the same calcium-binding proteins that we used to describe the chemoarchitecture of the PIL in rodent models, we were able to demonstrate the existence of a corresponding brain region in the human thalamus (Fig. 6A). Parvalbumin labeled the surrounding medial geniculate body but not the PIL, while calbindin-positive cells were abundant in the PIL region, indicating the existence of a chemoarchitecturally similar PIL in human. Further histological staining techniques also distinguished the PIL cytoarchitectonically from the surrounding brain regions (Fig. 6B).

**Figure 6.**
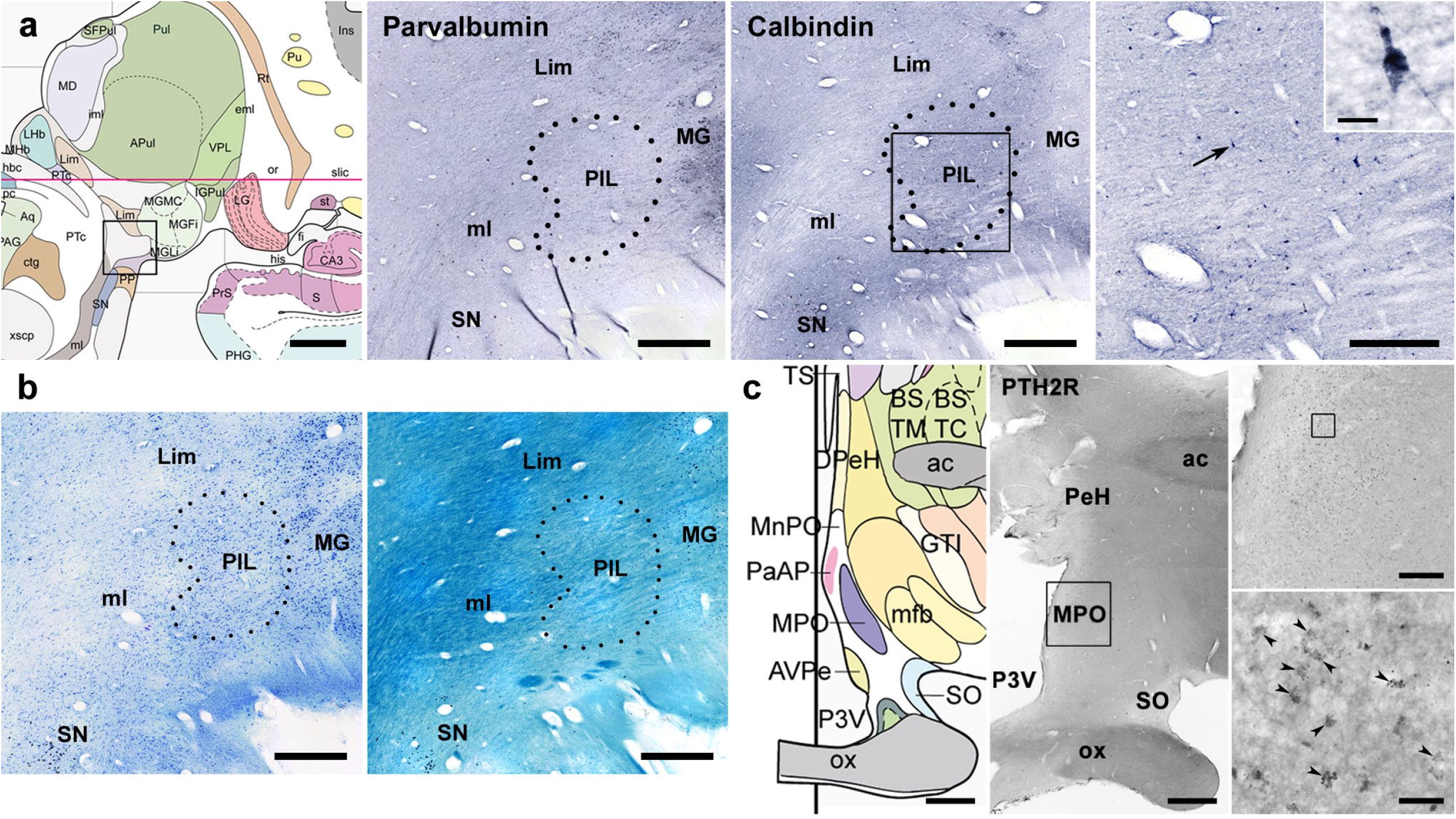
Elements of the PIL-MPOA pathway in human brain. **a,** The left panel shows a drawing of the PIL region (Mai, 2008), from which the framed area is enlarged in subsequent panels. Using calcium-binding proteins as markers, the location of the PIL was identified in the human thalamus. Parvalbumin intensely labelled the adjacent medial geniculate body but not the PIL, while calbindin appeared in the PIL but not in the MG. Scale bars: 500 μm (first image), 100 μm (second and third images), 40 μm and 20 μm (fourth image and its inset). **b,** Further histological staining using Nissl (left) and Luxol (right) also indicates the location of the PIL and how it is distinct from the surrounding brain regions. Scale bars: 100 μm. **c,** The expression of PTH2R, the receptor for PTH2, is demonstrated in the human medial preoptic area (indicated by black arrowheads). Scale bars: 2 mm (first two images), 300 μm (third upper image), 30 μm (third lower image).

In addition, the expression of PTH2R was observed in the human medial preoptic area, suggesting the existence of a PIL→MPOA projection in the human brain, as well (Fig. 6C).

## Discussion

The induction of PTH2 in response to social interaction generalizes previous findings on the elevated level of PTH2 in a mother rat in the presence of her litter (Cservenak et al., 2010). These data are also in line with a recent report that the level of the corresponding peptide, PTH2, in individual zebrafish is proportional to the total density of zebrafish in the surrounding environment (Anneser et al., 2020). In zebrafish, the cells expressing PTH2 were localized at the lateral edge of the *orthopedia homeobox a* (*otpa*) expression domain, which is known to control the specification of diencephalic neuroendocrine cells (Fernandes et al., 2013). The site of socially dependent expression of PTH2 in mammals - the PIL (Dobolyi et al., 2010) was also defined in human based on molecular anatomy (Nagalski et al., 2016). It corresponds to the area expressing PTH2 in the zebrafish based on the location of the cells. (For details of previous anatomical and functional investigations of the PIL area see SI Appendix, Table S2.)

While the results in the zebrafish study established how PTH2 expression is induced by conspecifics – using their lateral line organ to sense water vibrations elicited by their conspecifics, the actual function of PTH2 neurons remained unexplored. To address this question, we manipulated PIL neurons using chemogenetic tools in rats and found that the time of direct physical contacts between animals was increased by chemogenetic stimulation of the PIL. This behavior included both side-to-side contact as well as mounting on top of each other. Other types of social interactions, including sniffing or staying in the vicinity of each other when separated by walls were not affected. In addition, social recognition and preference for social novelty were also not changed. Therefore, we conclude that PIL neurons selectively promote social touch without affecting anxiety-like behaviors. In turn, the same increase in the duration of direct social contact was found when instead of chemogenetically stimulating all PIL neurons only those which were activated during a previous social interaction were stimulated, suggesting that promoting social touch is dependent on socially activated neurons. Since social touch is rewarding (Shamay-Tsoory and Eisenberger, 2021), it is possible that a subset of social contact experienced neurons in the PIL promotes further social contact to maintain social interaction with conspecifics.

The target regions of PIL neurons were similar to the location of PTH2 fibers in the brain, suggesting that PTH2 neurons are the major projection neurons of the PIL. Social interaction activated PTH2 neurons in the PIL as well as their target areas. Since not only social interaction, but also the chemogenetic stimulation of PIL neurons activated the target areas, we conclude that PTH2 neurons play an important role in conveying the effect of social interactions to a variety of forebrain areas implicated in social behavior.

PIL neurons projects to many brain regions including the prefrontal cortex, amygdala and hypothalamus. Here we show that the PIL projections to the MPOA are critical for mediation of social touch induced social behavior. Indeed, despite widespread brain activation in response to social interaction, only MPOA neurons were exclusively sensitive to direct physical contact, while olfactory, visual and auditory signals were insufficient to activate the MPOA. In support of the idea that PTH2 signaling contributed to the PIL→MPOA stimulated increase in direct social communication, PTH2-containing terminals closely appose socially activated PTH2 receptor expressing GABAergic neurons in the MPOA.

In conclusion, we discovered a novel subcortical pathway originating from PTH2 neurons in the PIL, which is exclusively activated by social touch and which in turn further promotes direct social interactions. The activity of this pathway may be crucial for a variety of behaviors including grooming, consolation and other nonsexual social touch, and also for reproductive behaviors, such as copulation, pair formation and maternal care in rodents (Tsuneoka and Funato, 2021) and probably in human. Thus, our findings can be relevant to pathophysiological mechanisms underlying avoidance of direct physical contacts/social touch with unfamiliar individuals and even with relatives, in patients afflicted with mental diseases. Therefore, the pathway should be investigated in the future in disorders where deficits in direct social interactions are found (Deschrijver et al., 2017), such as autism spectrum disorder (Ebert and Greenberg, 2013).

## Acknowledgements

Support was provided by NKFIH-4300-1/2017-NKP_17 (National Brain Research Program), OTKA K134221 research grant, Development and Innovation Fund and Eötvös Loránd University Thematic Excellence Programme 2020 (TKP2020-IKA-05) for AD; Gedeon Richter Plc. Centenary Foundation (Gyömrői út 19-21. Budapest, 1103), Excellence Program of the Semmelweis University, New National Excellence Program of the Ministry of Innovation and Technology, Doctoral Student Scholarship Program of the Co-operative Doctoral Program of the Ministry of Innovation and Technology financed from the National Research and EFOP-3.6.3-VEKOP-16-2017-00009 for DK; János Bolyai Research Scholarship of the Hungarian Academy of Sciences for MC; NIMH ZIC MH002963-05 for TU; 2017-1.2.1-NKP-2017-00002 for MP; the Ministry of Economy and Business (MINECO) under the framework of the ERA-NET Neuron Cofund and Ministerio Economia, Industria y Competitividad (Spain) and FEDER (grant RTI2018-101624-B-I00) for MTH; the German Research Foundation (DFG) grants GR 3619/8-1, GR 3619/13-1, GR 3619/15-1, GR 3619/16-1, Training Research Group (GRK) 2174 and SFB Consortium 1158-2 for VG. We also appreciate the technical assistance of Nikolett Hanák, Viktória Dellaszéga-Lábas and Szilvia Deák.

## Author contributions

DK participated in the design of the experiments, performing the behavioral and histological work, interpretation of the results and the literature, and writing of the manuscript. TL did some of the behavioral studies, MC, GP, and JB did some of the histological labeling and analysis, VC and IF performed the patch clamp experiment, DZ did the virus injections into mice, FD and MP collected the human brain tissue, TU contributed reagents and to editing the manuscript, LB produced vGATE viruses, MTH and VG participated in the design of the vGATE-based experiment, interpretation of the results, and correction of the manuscript. AD participated in the design of the experiments, interpretation of the results and the literature, and writing of the manuscript.

## STAR methods text

### Resource availability

#### Lead contact

Further information and requests for resources and reagents should be directed to and will be fulfilled by the lead contact, Arpad Dobolyi.

#### Materials Availability

This study did not generate new unique reagents.

#### Data and Code Availability

No custom code was used in the analysis of the data.

All data used in this manuscript are available upon request from the lead author.

### Experimental model and subject details

#### Animals

The Workplace Animal Welfare Committee of the National Scientific Ethical Committee on Animal Experimentation at Semmelweis and Eötvös Loránd Universities, Budapest, specifically approved this study (PE/EA/926-7/2021 and PE/EA/568-7/2020, respectively). Thus, the procedures involving rats and mice were carried out according to experimental protocols that meet the guidelines of the Animal Hygiene and Food Control Department, Ministry of Agriculture, Hungary (40/2013), which is in accordance with EU Directive 2010/63/EU for animal experiments. A total of 116 adult female rats (Wistar; Charles Rivers Laboratories, Hungary) were used (5 for labeling with PTH2 and calbindin, 20 for qRT-PCR study, 15 for social c-Fos study, 12 for chemogenetics / c-Fos study, 5 for chemogenetic stimulation of the PIL, 5 for chemogenetic inhibition of the PIL, 3 for viral injection into the SN, 10 for anterograde tracer injection into the PIL and SN, 28 for vGATE studies, 4 for double injections into the PIL and MPOA, 5 for cannula implantation into the MPOA for CNO administration, 4 for labeling with c-Fos, PTH2 and NeuN). In the behavioral experiments, vaginal smear tests were used to select diestrous rats for the testing. All of the rats were 90-120 days old when sacrificed. A total of 19 adult female mice were used in the study. There were 5 mice used for retrograde virus injection into the MPOA, 8 PTH2R-ZsGreen mice (offspring of PTH2R-Cre (generated by Ted B. Usdin, NIH) and GtROSA26Sor_CAG/ZsGreen1 mice (The Jackson Laboratory; stock number: 007906)) for labeling PTH2R and c-Fos, 3 VGAT-ZsGreen mice (offspring of VGAT-IRES-Cre and GtROSA26Sor_CAG/ZsGreen1 mice (The Jackson Laboratory; stock numbers: 016962 and 007906)) for labeling with VGAT and c-Fos/PTH2, 3 VGAT-ZsGreen mice for patch-clamp experiment). There were no behavioral or health issues found in VGAT-Cre and PTH2R-Cre mice crossed with reporter mice as we described previously (Dimen et al., 2021). All of the mice were 100-150 days old when sacrificed. Animals were kept under standard laboratory conditions with 12-h light, 12-h dark periods (lights on at 6.00 a.m.), and supplied with food and drinking water *ad libitum*. Pregnant and mother rats were individually housed. Mother rats who delivered fewer than 8 pups or whose pups died were excluded from the study. The number of pups was adjusted to 8 within 2 days of delivery. Animals were anaesthetized with an intramuscular injection of anesthetic mix containing 0.4 ml/300g body weight ketamine (50 mg/ml) and 0.2 ml/300g body weight xylazine (20 mg/ml) for surgery, perfusions and dissections.

#### Viral vectors

PIL stimulation or silencing was performed with the following viruses: pAAV-hSyn-hM3D(Gq)-mCherry (Addgene plasmid # 50474; http://n2t.net/addgene:50474; RRID: Addgene_50474) and pAAV-hSyn-hM4D(Gi)-mCherry (Addgene plasmid # 50475; http://n2t.net/addgene:50475; RRID: Addgene_50475), which were gifts from Bryan Roth. Activity-dependent tagging of PIL neurons was performed with a virus cocktail, which included the following viruses: 1. rAAV-(tetO)_7_-**P**_fos_-rtTA; 2. rAAV-**P**_tet_bi-Cre/YC3.60; 3. pAAV-hSyn-DIO-hM3D(Gq)-mCherry (a gift from Bryan Roth, Addgene plasmid # 44361; http://n2t.net/addgene:44361; RRID: Addgene_44361, Krashes et al., 2011) or pAAV-hSyn-DIO-hM4D(Gi)-mCherry (a gift from Bryan Roth, Addgene plasmid # 50475; http://n2t.net/addgene:50475; RRID: Addgene_50475) or pAAV-hSyn-DIO-mCherry (a gift from Bryan Roth, Addgene plasmid # 50459; http://n2t.net/addgene:50459; RRID: Addgene_50459), the first 2 viruses were the same as used in our previous study, Hasan et al., 2019.

For chemogenetic activation of PIL neurons projecting to the MPOA, first, a retrogradely spreading virus (AAVrg-Ef1a-mCherry-IRES-Cre, which was a gift from Karl Deisseroth (Addgene plasmid # 55632; http://n2t.net/addgene:55632; RRID: Addgene_55632)) (Fenno et al., 2014) was targeted to both sides of the MPOA. Then, three weeks later, a second virus (pAAV-hSyn-DIO-HA-hM3D(Gq)-IRES-mCitrine, a gift from Bryan Roth (Addgene plasmid # 50454; http://n2t.net/addgene:50454; RRID: Addgene_50454) was injected into the PIL in both sides.

### Method details

#### Microdissection of brain tissue samples

For qRT-PCR, brains were dissected from 10 female adult rats kept in social environment (3 animals/cage) and 10 female adults isolated for two weeks prior to dissection. Immediately after the dissections of the brain, the PIL was microdissected as described previously in mother rats (Cservenak et al., 2010). 1 mm thick coronal brain sections were cut from the freshly dissected brains. The area containing the PIL was subsequently microdissected from the appropriate coronal section with a circular micropunch needle of 1 mm diameter. The dissected tissue samples were quickly frozen in Eppendorf tubes on dry ice, and stored at −80°C until further processing.

#### Real-time PCR

Real-time qRT-PCR was carried out as described previously (Dobolyi, 2009). Briefly, total RNA was isolated from frozen PIL tissue samples using TRIzol reagent (Invitrogen, Carlsbad, CA, USA) as lysis buffer combined with RNeasy Mini kit (Qiagen, Germany) following the manufacturer’s instructions. The concentration of RNA was adjusted to 500 ng/µL, and it was treated with Amplification Grade DNase I (Invitrogen). Then, cDNA was synthesized using SupersciptII (Invitrogen) as described in the kit protocol. The cDNA was subsequently diluted (10x), and 2.5 µL of the resulting cDNA was used as template in multiplex PCR performed in duplicates using dual-fluorescence labeled TaqMan probes for *PTH2* (GenBank Accession Number (Rattus norvegicus): NM_001109144.1; 6-FAM-CGCTAGCTGACGACGCGGCCT-TAMRA), and glyceraldehyde-3-phosphatedehydrogenase (*GAPDH*; GenBank Accession Number (Rattus norvegicus): NM_017008.4; JOE-ATGGCCTTCCGTGTTCCTACCCCC-TAMRA) as the housekeeping gene as described previously (Cservenak et al., 2010). The primers for *PTH2* (CTGCCTCAGGTGTTGCCCT and TGTAAGAGTCCAGCCAGCGG) were used at 300 nM, whereas the primers for *GAPDH* (CTGAACGGGAAGCTCACTGG and GGCATGTCAGATCCACAAC) were used at 150 nM concentration. The PCR reactions were performed with iTaq DNA polymerase (Bio-Rad Laboratories, Hercules, CA, USA) in total volumes of 12.5 μl under the following conditions: 95°C for 3 min, followed by 35 cycles of 95°C for 0.5 min, 60°C for 0.5 min, and 72°C for 1 min. Cycle threshold (Ct) values were obtained from the linear region of baseline adjusted amplification curves and duplicates were averaged. The data were expressed as the ratio of the housekeeping gene GAPDH using the following formula: log(Ct(GAPDH)-Ct(PTH2)). Statistical analyses were performed by unpaired t-test for comparisons of the two groups.

### Histology

#### Animal brain tissue collection and sectioning

Animals were deeply anesthetized and perfused transcardially with 150 ml of saline followed by 300 ml of ice-cold 4% paraformaldehyde prepared in 0.1 M phosphate buffer at pH = 7.4 (PB) for rats and with 15 ml of saline followed by 30 ml of ice-cold 4% paraformaldehyde prepared in 0.1 M PB for mice. Brains were removed and postfixed in 4% paraformaldehyde for 24 h, and then transferred to PB containing 20% sucrose for 2 days. Serial coronal sections were cut at 40 μm on a sliding microtome and sections were collected in PB containing 0.05% sodium-azide and stored at 4 °C.

#### Immunolabeling

Every fifth free-floating brain section in rats and every third in mice were used for each labeling. First, the sections were pretreated in PB containing 0.3% hydrogen peroxide for 15 min to eliminate endogenous peroxidase activity, then in PB containing 0.5% Triton X-100 and 3% bovine serum albumin for 1 h for antibody penetration and reduction of nonspecific labeling, respectively. Then, primary antisera (SI Appendix, Table S3) were applied for 24 h at room temperature. This was followed by incubation of the sections in biotinylated or fluorescent secondary antibodies (Jackson ImmunoResearch, SI Appendix, Table S3) for 1 h. For non-fluorescent single labeling and for tyramide amplification fluorescent labeling, the sections were incubated in avidin-biotin-peroxidase complex (ABC, 1:500; Vector Laboratories) for 1 h. Subsequently, sections were treated with 0.02% 3,3’-diaminobenzidine (DAB; Vector Laboratories), 0.08% nickel (II) sulfate and 0.003% hydrogen peroxide in Tris hydrochloride buffer (0.1 M, pH = 8.0) for 6 min for non-fluorescent immunoperoxidase technique, and with FITC-tyramide (1:5,000) and H_2_O_2_ for tyramide amplification immunofluorescence. For double labeling, another primary antibody (SI Appendix, Table S3), produced in a different species as the first antibody, was added at room temperature overnight. Sections were incubated in secondary antibodies produced in donkey and labeled with Alexa dyes (1:500; Jackson ImmunoResearch) for 2 h. Finally, the sections were mounted, dehydrated and coverslipped with DePeX Mounting Medium (Sigma) for non-fluorescent, and Aqua-Poly/Mount (VWR) for fluorescent labeling.

### Anterograde tracing with biotinylated dextran amine

#### Injection of anterograde tracer into the PIL

The anterograde tracer biotinylated dextran amine (BDA, 10,000 MW; Molecular Probes) was targeted to the PIL (n=5). Some of the misplaced injections were used as controls (n=5). For stereotaxic injections, rats were positioned in a stereotaxic apparatus with the incisor bar set at −3.3 mm. Holes of about 1 mm diameter were drilled into the skull above the target coordinates. Glass micropipettes of 15–20 μm internal diameter were filled with 10% BDA dissolved in PB and lowered to the following stereotaxic coordinates (Paxinos and Watson, 2007): AP = −5.2 mm from bregma, ML = +2.6 mm from the midline, DV = −6.8 mm from the surface of the dura mater. Once the pipette was in place, the BDA was injected by iontophoresis by using a constant current source (51413 Precision Current Source, Stoelting, Wood Dale, IL) that delivered a current of +6 μA, which pulsed for 7 sec on and 7 sec off for 15 min. Then the pipette was left in place for 10 min with no current and withdrawn under negative current. After tracer injections, the animals were allowed to survive for 7 days. Visualization of BDA was done as described above for Ni-DAB visualization.

### Social c-Fos activation study

#### Social interaction experiment

Female rat littermates (n=15) kept together (housed 2 per cage) were used in the study. The 2 rats were isolated for 22 h to reduce basal c-Fos activation. Then, the rats were reunited again in the cage they had cohabited. 7 rats could freely interact with their cagemate while 4 rats were isolated from their partner with bars, so they could see, hear and smell each other but could not directly interact. Control females (n=4) continued to be kept isolated. All animals were sacrificed 24 h after the beginning of isolation, i.e. 2 h after reunion of the socially interacting group. Animals were perfused transcardially as described above, and processed for c-Fos immunohistochemistry.

#### c-Fos immunohistochemistry

The procedure was performed as described above for immunolabeling using rabbit anti-c-Fos primary antiserum (1:3000; Santa Cruz Biotechnology) and Ni-DAB visualization.

#### Analysis of c-Fos immunolabeling

The sections containing the greatest number of c-Fos-ir neurons in the investigated brain regions (ILC, LS, MPOA, PVN, MeA, DMH, VLPAG) were selected from each of the 15 animals, and an image was taken of them using 10x objective. The total number of c-Fos-ir neurons in the selected brain area was counted using ImageJ 1.53e program (NIH, Bethesda, MD, Abràmoff et al., 2004) as described previously (Cservenak et al., 2017). Briefly, the same algorithm, based on intensity, size, and circularity thresholding was used for all images to select the Fos-ir nuclei, whose number was counted automatically by the program. Spots were selected whose brightness intensity value was <150. The number of c-Fos–containing cells was defined as the number of selected spots counted in a size ranging from 20 to 100 pixels, and a circularity factor between 0.5 and 1.0. Statistical analyses were performed using Prism 9 for Windows (GraphPad Software, Inc., La Jolla, CA). The number of c-Fos-ir neurons in the 3 groups (after social interaction, social interaction without direct contact, isolated control) was compared using two-way ANOVA.

### Viral vector-based chemogenetics

#### Chemogenetic activation or silencing of the PIL

pAAV-hSyn-hM3D(Gq)-mCherry (Addgene plasmid # 50474; http://n2t.net/addgene:50474; RRID: Addgene_50474) and pAAV-hSyn-hM4D(Gi)-mCherry (Addgene plasmid # 50475; http://n2t.net/addgene:50475; RRID: Addgene_50475), were injected to the PIL (n=5 for each group). Some of the misplaced injections were used as controls in tract tracing (n=3). For stereotaxic injections, rats were positioned in a stereotaxic apparatus with the incisor bar set at −3.3 mm. Holes of about 1 mm diameter were drilled into the skull above the target coordinates. A Hamilton pipette (volume 1 μl) was filled with the solution containing the virus and lowered to the following stereotaxic coordinates (Paxinos and Watson, 2007): AP = −5.2 mm from bregma, ML = ±2.6 mm from the midline, DV = −6.8 mm from the surface of the dura mater. Once the pipette was in place, 100 nl virus was pressure injected into the PIL (10 nl/min), then the pipette was left in place for 10 min. Following the slow withdrawal of the pipette, we repeated the same protocol on the other side of the brain. After the injections, the animals were isolated for two weeks for recovery, then they were reunited with their familiar cagemate for one week before the behavior tests took place.

#### Activity-dependent tagging: Chemogenetic manipulation of PIL neurons activated upon social encounter

The protocol for bilateral injections of the vGATE viral cocktail into the PIL was as described above. The viral cocktail included the following viruses: 1. rAAV-(tetO)_7_-**P**_fos_-rtTA; 2. rAAV-**P**_tet_bi-Cre/YC3.60; 3. pAAV-hSyn-DIO-hM3D(Gq)-mCherry (excitatory, n=12), or pAAV-hSyn-DIO-hM4D(Gi)-mCherry (inhibitory, n=8) or pAAV-hSyn-DIO-mCherry (control, n=8). 600 nl of single injection contained the following: 1^st^ virus: 225 nl. 2^nd^ virus: 75 nl, 3^rd^ virus: 300 nl (3/1/4 ratio). After two weeks of recovery, the animals were treated with i.p. doxycycline hyclate injection (5 mg/kg bw dissolved in the mixture of 0.01M PBS and 0.9% NaCl, 2 : 1 ratio), then, the following day, they were reunited with one of their cagemates for 2 hours for expression of DREADD in neurons that are c-Fos-positive in the PIL in response to social interaction. The behavior tests started 10 days later in analogy to Hasan *et al*., 2019.

#### Chemogenetic activation of PIL neurons projecting to the MPOA

First, a retrogradely spreading virus (AAVrg-Ef1a-mCherry-IRES-Cre, which was a gift from Karl Deisseroth (Addgene plasmid # 55632; http://n2t.net/addgene:55632; RRID: Addgene_55632)) (Fenno et al., 2014) was targeted to both sides of the MPOA. Then, three weeks later, a second virus (pAAV-hSyn-DIO-HA-hM3D(Gq)-IRES-mCitrine, a gift from Bryan Roth (Addgene plasmid # 50454; http://n2t.net/addgene:50454; RRID: Addgene_50454) was injected into the PIL in both sides (n=4). The viral injections were carried out as described above. The stereotaxic coordinates of the MPOA was the following (Paxinos and Watson, 2007): AP = −0.5 mm from bregma, ML = ±0.6 mm from the midline, DV = −6.7 and −7.7 mm from the surface of the dura mater. The MPOA was targeted in two ventral coordinates, 100 nl of virus was injected at both levels.

#### Chemogenetic activation of axonal terminals of PIL neurons in the MPOA

Implantation of cannulae into the MPOA was performed immediately after the injection of the excitatory virus (pAAV-hSyn-hM3D(Gq)-mCherry, see above) as described above. In rats (n=5) already anaesthetized with 0.2 ml xylazine and 0.4 ml ketamine and fixed in a stereotaxic apparatus, a hole of about 1 mm diameter was drilled into the skull above at the following coordinates (Paxinos and Watson, 2007): AP = −0.5 mm from bregma, ML = ±3.0 mm from the midline. Cannulae (ALZET Brain Infusion Kit 2, Durect™) were inserted into the MPOA (V = 7.2 mm) with an inclination angle of 16.5° and fixed to the skull with cranioplastic cement. Subsequently, the same protocol was repeated on the other side. After the operation, Tardomyocel® comp. III antibiotics (0.1 ml/kg bw) was given s.c. to the animals for 5 days to prevent infections.

### Chemogenetics-based behavioral tests

#### Clozapine-N-oxide (CNO) injection

All behavioral tests were performed on adult female rats to avoid aggressive components of social interactions that often occur in males and in analogy to our previous study (Tang et al., 2020). 1.5 h before the behavioral experiments, the animals were isolated from their cagemates, which was followed by the intraperitoneal administration of CNO or vehicle. On the first day of the experimental session, we used control vehicle injection (5% DMSO dissolved in distilled water, 1 ml/kg bw). One or two days later, CNO was administered i.p. (0.3 mg/kg bw CNO, dissolved in 5% DMSO dissolved in distilled water, 1 ml/kg bw). This was followed by another control injection three or four days later in certain experiments (vGATE experiments, chemogenetic stimulations of the PIL→MPOA pathway).

#### Test of social interaction

During the experiment, the animals were placed in an open field arena (40 x 80 cm) where the subject and the stimulus rat could freely interact with each other and half an hour of footage was recorded. Different social behavior elements were distinguished, which were categorized as direct (physical) social contact (including “mounting”, “being mounted”, “side-to-side contact”) and social sniffing (including “body sniffing”, “head-to-head contact”) in analogy to Tang et al., 2020. In another experiment, direct contact between the animals was not allowed. Instead, the stimulus rat was placed in a small cage with wall made of bars, so they could just smell, hear and see each other while 10 min of footage was recorded.

The duration of the different types of performed behaviors were measured by using Solomon Coder software (https://solomon.andraspeter.com/). The behavior of the animals in the experiment excluding the physical contact was analyzed with SMART Video Tracking software v3.0 (Panlab Harvard Apparatus). The duration of the behavioral elements was compared between the groups using Student’s t test.

#### Social novelty test

The experiment measuring the preference of social novelty was conducted in the Three Chambers Sociability Apparatus (120 x 80 cm, including three 40 x 80 cm chambers connected with doors). 1.5 hour after the administration of drug (control or CNO injection, see above), the subject was placed in the middle chamber. Each side chamber contained a small cage with wall made of bars, in one of them the familiar cagemate of the subject, in the other one an unfamiliar animal were placed. During the 10 min of the experiment, the subject could freely move between the three chambers. The behavior of the animal was analyzed with SMART Video Tracking software v3.0 (Panlab Harvard Apparatus). The duration of the time spent in the chambers was compared between the groups using Student’s t test.

#### Depression and anxiety-like behavior tests

These behavioral tests cannot be repeated in the same animal. Therefore, the subjects in each group were split into two: half of them were administered a control injection, the other half received CNO injection. The depression-like behavior was measured with the forced swim test (FST). The animals were placed in a container containing water (24 °C), from which they could not escape. The time spent climbing and floating were measured for 6 min. The videotapes were analyzed with Solomon Coder software (https://solomon.andraspeter.com/).

The anxiety-like behavior was measured with the elevated plus maze test (EPM), where the animals were placed in a maze, which was 60 cm above the ground, and composed of two open and two closed arms. The animals were placed in the center of the maze and could freely move in all the four arms. The time spent in the open and closed arms was measured for 10 min. The trajectory of the animals was analyzed with SMART Video Tracking software v3.0 (Panlab Harvard Apparatus). The duration of the time spent in the open/closed arms was compared between the groups using the Student’s t test.

### Induction of c-Fos expression driven by chemogenetic stimulation

Female rats (n=12) were injected with excitatory virus (pAAV-hSyn-hM3D(Gq)-mCherry) into the PIL as described above. Three weeks later, 6 animals received control i.p. injection (5% DMSO dissolved in distilled water, 1 ml/kg bw) and 6 animals were injected i.p. with CNO (0.3 mg/kg bw CNO, dissolved in 5% DMSO dissolved in distilled water, 1 ml/kg bw). All animals were sacrificed 2 h after the drug administration. Animals were perfused transcardially and processed for c-Fos immunohistochemistry as described above. The sections containing the greatest number of c-Fos-ir neurons in the investigated brain regions (PIL, ILC, LS, MPOA, PVN, MeA, DMH, VLPAG) were selected from each of the 12 animals and analyzed as described above for social c-Fos study. The number of c-Fos-ir neurons in the two groups was compared using two-way ANOVA.

### Immunohistochemical characterisation of neurons in the PIL→MPOA pathway

#### Double labeling of PTH2 with calbindin or parvalbumin

Every fifth free-floating brain section was immunolabeled with tyramide amplification fluorescent immunolabeling using an affinity-purified anti-PTH2 antiserum (1:3,000) validated previously (Dobolyi et al., 2002; Wang et al., 2006). Then, mouse anti-calbindin D-28k (1:1,500; Sigma) or mouse anti-parvalbumin antiserum (1:2,500; Sigma) was applied at room temperature overnight. Following the application of the primary antiserum, sections were incubated in donkey Alexa Fluor 594 anti-mouse secondary antibody (1:500; Jackson ImmunoResearch) for 2 h.

#### Single labeling of mCherry

The injection sites of viruses as well as the projection pattern of the infected neurons was studied using immunoperoxidase technique with the visualization of mCherry, for which a chicken anti-mCherry antiserum (1:1,000; Abcam) was used with Ni-DAB method.

For fiber density analysis of PIL projections, we used the machine learning based pixel classification feature of the free bioimage analysis software QuPath v0.2.3 (Queen’s University Belfast and University of Edinburg) (Bankhead et al., 2017). We determined the area density of the anterogradely labeled nerve fibers (mCherry immunolabeled fiber area/total area) in socially implicated brain regions.

#### Double labeling of mCherry with c-Fos, calbindin, parvalbumin and mCitrine

First, rabbit anti-c-Fos primary antiserum (1:750; Santa Cruz Biotechnology) or mouse anti-calbindin D-28k antiserum (1:1,500; Sigma) or mouse anti-parvalbumin antiserum (1:2,500; Sigma) or goat anti-GFP antiserum (1:1,000; Abcam) for mCitrine labeling was applied for 24 h at room temperature. This was followed by incubation of the sections in biotinylated donkey anti-rabbit/mouse/goat secondary antibodies (1:1,000; Jackson ImmunoResearch). Subsequently, sections were treated with FITC-tyramide (1:5,000) and H_2_O_2_ in Tris hydrochloride buffer (0.1 M, pH = 8.0) for 6 min. Then, chicken anti-mCherry antiserum (1:1,000; Abcam) was applied at room temperature overnight. Following application of the primary antiserum, sections were incubated in donkey Alexa Fluor 594 anti-chicken secondary antibody (1:500; Jackson ImmunoResearch) for 2 h.

#### Triple labeling of c-Fos, PTH2 and NeuN

To visualize the neuronal activation upon social encounter in the MPOA, the animals (n=4) kept in pairs were isolated for 22 h to reduce basal c-Fos activation. Then, the rats were reunited again in the cage they had cohabited for 2 h. Animals were perfused transcardially as described above, and processed for immunohistochemistry. First, rabbit anti-PTH2 primary antiserum (1:3,000) was applied and visualized with FITC-tyramide. Then, rabbit anti-c-Fos primary antiserum (1:750; Santa Cruz Biotechnology) was applied and visualized with donkey Alexa Fluor 594 anti-rabbit secondary antibody (1:500; Jackson ImmunoResearch) for 2 h. Finally, mouse anti-NeuN primary antiserum (1:500; Merck Millipore) was applied at room temperature overnight. Following the application of the primary antiserum, sections were incubated in donkey Cy5 anti-mouse secondary antibody (1:300; Jackson ImmunoResearch) for 2 h.

#### Labeling of c-Fos in PTH2R-ZsGreen mice

To visualize the neuronal activation of PTH2R-positive neurons upon social encounter in the MPOA, social c-Fos study was conducted as described above in PTH2R-ZsGreen mice (n=4 for social group, n=4 for isolated control). Then, the brains were processed for c-Fos immunohistochemistry as described above using visualization with Alexa Fluor 594. Then, 4′,6-diamidino-2-phenylindole (DAPI, 1:10,000) was applied at room temperature for 5 min. The total number of double labeled neurons with c-Fos and PTH2R in the MPOA in the two group was counted using QuPath v0.2.3 (Queen’s University Belfast and University of Edinburg) (Bankhead et al., 2017).

#### Labeling of c-Fos/PTH2 in VGAT-ZsGreen mice

To visualize the neuronal activation of VGAT-positive neurons upon social encounter in the MPOA, a social c-Fos study was conducted in VGAT-ZsGreen mice as described above. The sections were processed for c-Fos and PTH2 immunolabeling using rabbit anti-c-Fos (1:500; Santa Cruz Biotechnology) or rabbit anti-PTH2 primary antisera (1:3,000).

### Histology in the human brain

#### Human brain tissue samples

Human brain (n=2) samples were collected in accordance with the Ethical Rules for Using Human Tissues for Medical Research in Hungary (HM 34/1999) and the Code of Ethics of the World Medical Association (Declaration of Helsinki). Tissue samples were taken during brain autopsy at the Department of Forensic Medicine of Semmelweis University in the framework of the Human Brain Tissue Bank (HBTB), Budapest. The activity of the HBTB has been authorized by the Committee of Science and Research Ethic of the Ministry of Health Hungary (ETT TUKEB: 189/KO/02.6008/2002/ETT) and the Semmelweis University Regional Committee of Science and Research Ethic (No. 32/1992/TUKEB), including removal, collecting, storing and international transportation of human brain tissue samples and applying them for research. Prior written informed consent was obtained from the next of kin, which included the request to consult the medical chart and to conduct neurochemical analyses. The study reported in the manuscript was performed according to protocol approved by the Committee of Science and Research Ethics, Semmelweis University (TUKEB 189/2015). The medical history of the subjects (62 and 79 years old females died in sepsis and myocardial infarction, respectively) was obtained from clinical records, interviews with family members and relatives, as well as from pathological and neuropathological reports. All personal identifiers had been removed and samples were coded before the analyses of tissue. Brains were dissected from the skull with a *post-mortem* delay of 24-48 h. Brains were cut into 5–10 mm thick coronal slices and immersion fixed in 4% paraformaldehyde in 0.1 M phosphate buffer (PB) for 6–10 days.

For histological labeling, coronally oriented tissue blocks of about 10×10×20 mm containing the preoptic area and the PIL, respectively, were sectioned in the coronal plane. Tissue blocks containing the preoptic area (from 1 mm rostral to the anterior commissure to the caudal end of the preoptic area) and the posterior thalamic region containing the PIL were sectioned. Two days before sectioning, the tissue blocks were transferred to PB for 2 days to remove the excess paraformaldehyde. Subsequently, the blocks were placed in PB containing 20% sucrose for 3 days for cryoprotection. Then, the blocks were frozen, and cut into 50 µm thick serial coronal sections on a sliding microtome.

#### Histological labeling in human brain sections

For cresyl-violet staining, sections were mounted consecutively on gelatin-coated slides and dried. Sections were stained in 0.1% cresyl-violet dissolved in PB and then differentiated in 96% ethanol containing acetic acid.

For Luxol fast blue staining, myelinated fibers were visualized first with the sulphonated copper phthalocyanine Luxol Fast Blue with a modification of the Kluver–Barrera method as described before (McIlmoyl, 1965). Briefly, the sections were stained in 0.1% Luxol fast blue dissolved in 96% ethanol containing 0.05% acetic acid, and differentiated in 0.05% lithium-chloride followed by 70% ethanol. Subsequently, sections were stained in 0.1% cresyl-violet. At the end, sections were dehydrated and coverslipped as described above.

#### Immunolabeling in human brain sections

Immunolabeling was performed as described above on every 10th section. The free-floating sections were pretreated with 3% bovine serum albumin in PB containing 0.5% Triton X-100 for 30 min at room temperature. The sections were then placed in rabbit anti-PTH2R (1:20,000), mouse anti-calbindin D-28k (1:3,000; Sigma) or mouse anti-parvalbumin (1:5,000; Sigma) primary antiserum for 48 h at room temperature. The custom made anti-PTH2R antiserum was previously used and characterized (Wang et al., 2000). Following the incubation in primary antisera, the sections were placed in biotinylated anti-rabbit/mouse secondary antibody (1:600; Vector Laboratories) for 2 h followed by incubation in a solution containing avidin–biotin– peroxidase complex (ABC, 1:300; Vector Laboratories) for 2 h. The sections were then treated with fluorescein isothiocyanate (FITC)-tyramide (1:8,000) and H_2_O_2_ (0.003%) in Tris hydrochloride buffer (0.05 M, pH 8.2) for 6 min. The sections were then mounted, dried, and coverslipped in mounting medium.

### Microscopy and image processing

Sections were examined using an Olympus BX60 light microscope equipped with fluorescent epi-illumination. Images were captured at 2048 X 2048 pixel resolution with a SPOT Xplorer digital CCD camera (Diagnostic Instruments, Sterling Heights, MI) using a 4-40 X objectives. Confocal images were acquired with a Zeiss LSM 780 confocal Microscope System using 40-63 X objectives at an optical thickness of 1 µm for counting varicosities and 3 µm for counting labeled cell bodies.

For demonstration purposes, contrast and sharpness of the images were adjusted using the “levels” and “sharpness” commands in Adobe Photoshop CS 9.0. Full resolution of the images was maintained until the final versions were adjusted to a resolution of 300 dpi.

### In vitro electrophysiological measurement

#### Brain slice preparation

Brain slice preparation from adult VGAT-IRES-Cre/Gt (ROSA)26Sor_CAG/ZsGreen1 female mice (n=3) was carried out as described previously (Farkas et al., 2010) with slight modification. Briefly, after deep isoflurane anaesthesia, the animal was decapitated, the brain was removed from the skull and immersed in ice-cold low-Na cutting solution, continuously bubbled with carbogen, a mixture of 95% O_2_ and 5% CO_2_. The cutting solution contained the following (in mM): saccharose 205, KCl 2.5, NaHCO_3_ 26, MgCl_2_ 5, NaH_2_PO_4_ 1.25, CaCl_2_ 1, glucose 10. Hypothalamic blocks containing the preoptic area were dissected, and 220 μm-thick coronal slices were prepared from the medial preoptic area (MPOA) with a VT-1000S vibratome (Leica Microsystems, Wetzlar, Germany) in the ice-cold low-Na oxygenated cutting solution. The slices were transferred into artificial cerebrospinal fluid (aCSF) (in mM): NaCl 130, KCl 3.5, NaHCO_3_ 26, MgSO_4_ 1.2, NaH_2_PO_4_ 1.25, CaCl_2_ 2.5, glucose 10 bubbled with carbogen and left in it for 1 hour to equilibrate. Equilibration started at 33°C and it was let to cool down to room temperature.

#### Whole cell patch clamp recording

Recordings were carried out in oxygenated aCSF at 33°C. Axopatch-200B patch-clamp amplifier, Digidata-1322A data acquisition system, and pCLAMP 10.4 software (Molecular Devices Co., Silicon Valley, CA, USA) were used for recording. VGAT-ZsGreen neurons were visualized with a BX51WI IR-DIC microscope (Olympus Co., Tokyo, Japan) after identifying them by their fluorescent signal. The patch electrodes (OD = 1.5 mm, thin wall; WPI, Worcester, MA, USA) were pulled with a Flaming-Brown P-97 puller (Sutter Instrument Co., Novato, CA, USA). The intracellular pipette solution contained (in mM): K-gluconate 130, KCl 10, NaCl 10, HEPES 10, MgCl_2_ 0.1, EGTA 1, Mg-ATP 4, Na-GTP 0.3. pH=7.2-7.3 with KOH. Osmolarity was adjusted to 300 mOsm with D-sorbitol. Electrode resistance was 2–3 MΩ. Firing activity of VGAT-ZsGreen neurons was recorded in whole-cell current clamp mode. In order to evoke action potentials even in silent neurons (approx. 30% of the neurons measured) +10 pA holding current was applied during the recordings. Measurements started with a control recording, then PTH2 (8 μM-32 μM, Tocris, UK) was pipetted into the aCSF-filled measurement chamber containing the brain slice in a single bolus and the recording continued. A previously identified selective and potent PTH2 receptor antagonist, HYWH-TIP39 (Kuo and Usdin, 2007) was pipetted into the recording chamber just before adding PTH2.

Input resistance and series resistance were monitored at the beginning and at the end of each experiment from the membrane current response to a 20 ms, 5 mV hyperpolarizing voltage step to monitor recording quality. Baseline voltage was also monitored before and after the experiments. All recordings with input resistances (*R*in) < 0.5 GΩ, and series resistances (*R*s) > 20 MΩ, or unstable baseline voltage were rejected.

#### Analysis of patch clamp experiment

Recordings were stored and analyzed off-line. Event detection was performed using the Clampfit module of the PClamp 10.4 software (Molecular Devices Co., Silicon Valley, CA, USA). Firing rate was calculated as number of action potentials (APs) divided by the length of the corresponding time period (1-1.5-5 min). Area-under-curve values were calculated from these data. Group data were expressed as mean ± standard error of mean (s.e.m). Repeated measures ANOVA was applied to examine whether firing rate change of each group was significant (*p* < 0.050).

### Statistics

All statistical calculations were carried out using GraphPad Prism (GraphPad Software, LLC, released 2020, version 9.0.0.). Data were first tested with Shapiro-Wilk test for normality. If the data were normally distributed, we used Student’s t-test for two groups. When the comparison was made between two different groups, we used unpaired Student’s t-test (for example in case of the qRT-PCR study or analysis of depression and anxiety-like behaviors). When the groups were related to each other (for example in case of self-control measurement in most of the social behavior tests) we used paired Student’s t-test. Otherwise, if the data came from a non-normal distribution, we performed Wilcoxon matched-pairs signed rank test. In case of the comparison between more groups (and the data were normally distributed), two-way ANOVA followed by Šídák’s multiple comparisons tests or repeated measures ANOVA were performed. We list the *p* value in all cases. Statistical analyses were considered significant for *p* < 0.050 (0.010 < \**p* < 0.050, 0.001 < \*\**p* < 0.010, \*\*\**p* < 0.001). All values are expressed as the mean ± s.e.m.

## Supplemental item titles and legends

**Figure S1.**
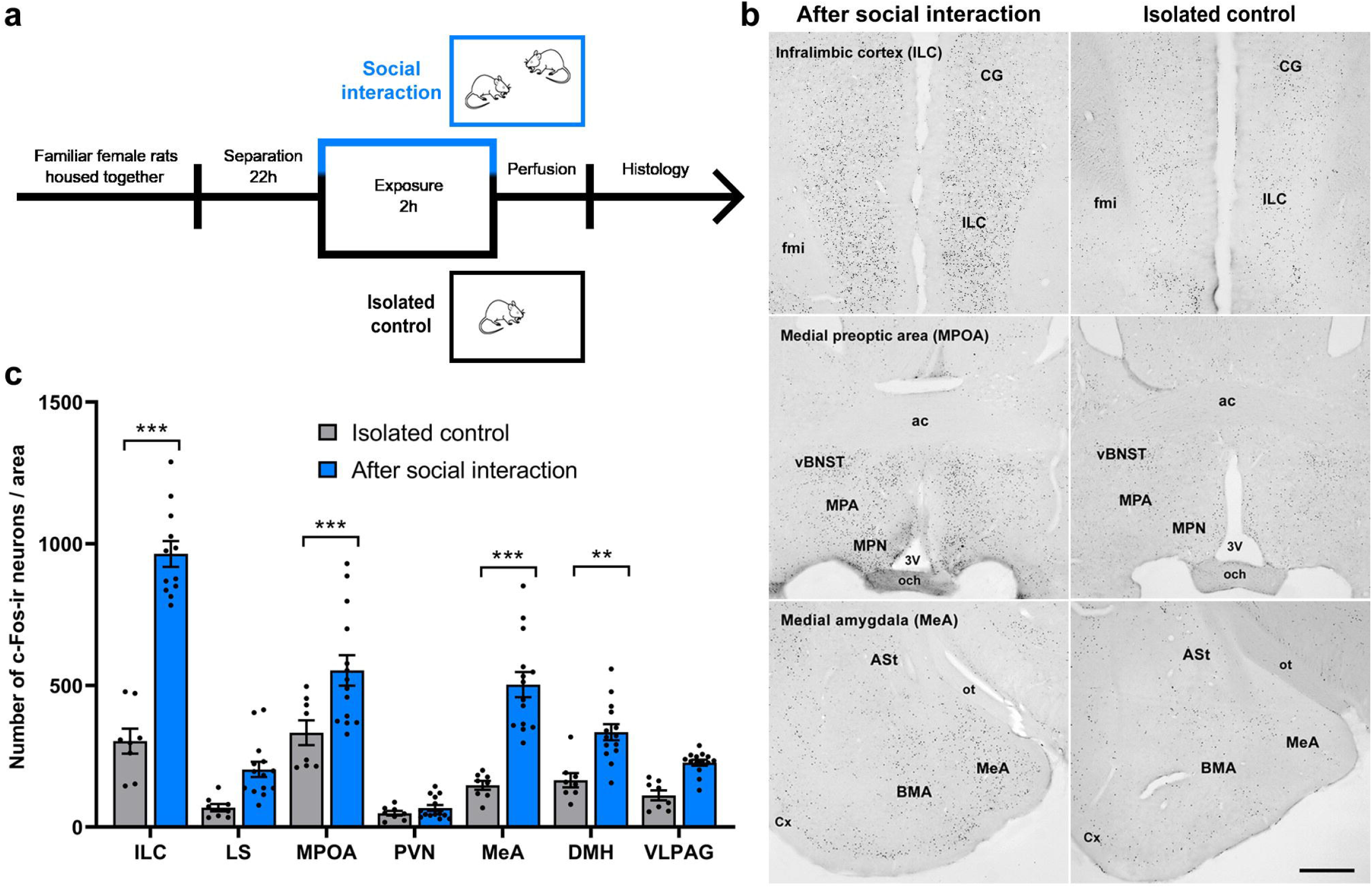
Activation of target areas of the PIL upon social encounter. **a,** Experimental protocol for studying neuronal activation in response to social interaction. **b,** Images of the c-Fos immunolabeling demonstrate the elevated c-Fos activation in ILC, MPOA and the amygdala upon social encounter. In isolated animals, only a low density of c-Fos-labeled neurons were present in these regions. Scale bar: 750 μm. **c,** Comparison of the c-Fos activation of seven socially implicated brain regions. For isolated control group: n=8, for social interaction: n=14, two-way ANOVA followed by Šídák’s multiple comparisons tests, for ILC: *p* < 0.001, for LS: *p* = 0.056, for MPOA: *p* < 0.001, for PVN: *p* = 0.99, for MeA: *p* < 0.001, for DMH: *p* = 0.0067, for VLPAG: *p* = 0.15; 0.001 < \*\**p* < 0.010 and \*\*\**p* < 0.001. Values are mean ± s.e.m.

**Figure S2.**
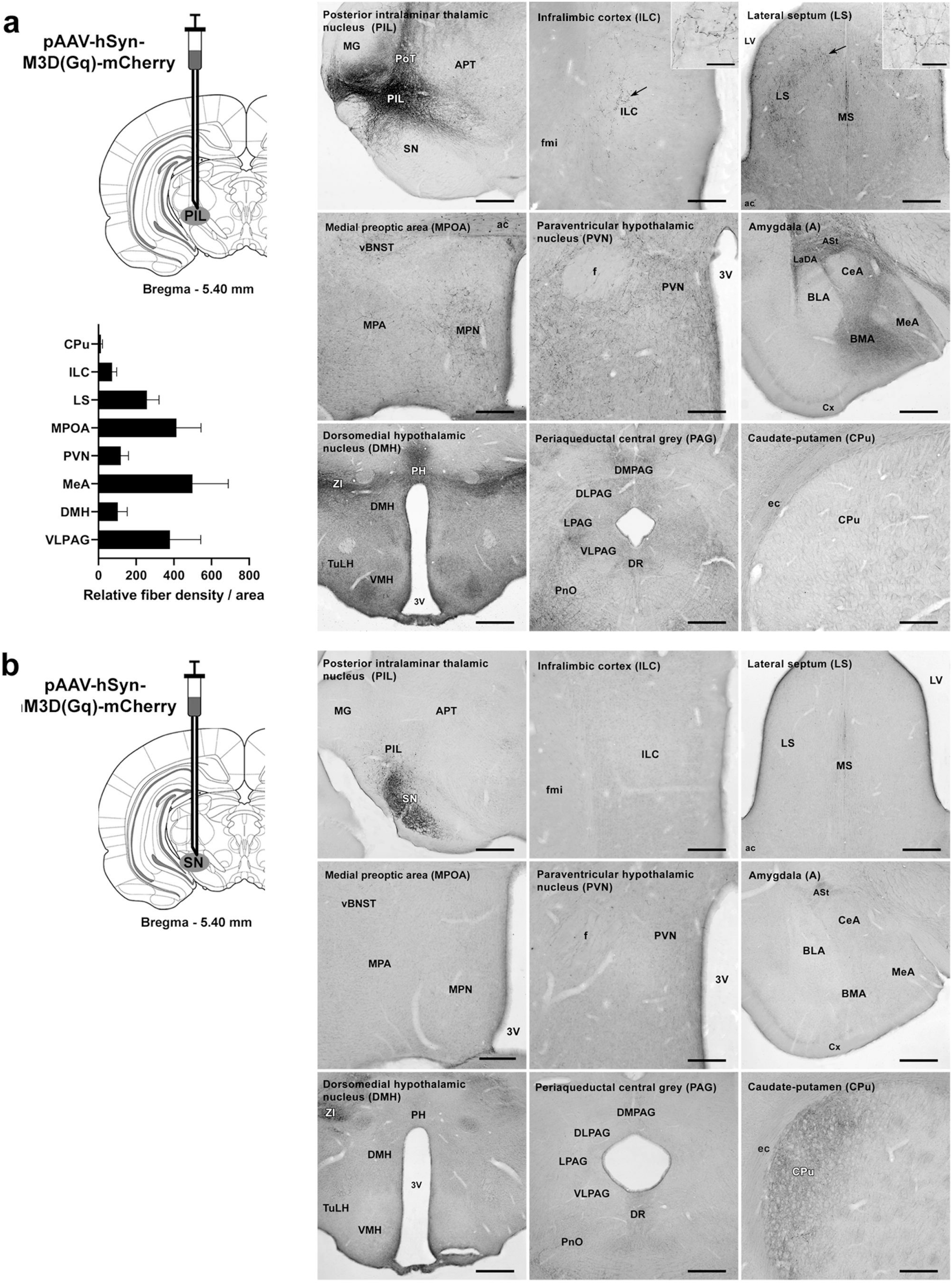
The anterograde spread of the virus injected into the PIL to socially implicated brain regions. **a,** The virus (pAAV-hSyn-hM3D(Gq)-mCherry) was injected into the PIL followed by mCherry immunolabeling to visualize DREADD-containing cells and fibers. Among the socially implicated brain regions, the highest density of the mCherry-positive fibers was found in the MPOA and the MeA. mCherry immunolabeled area / total area of the annotation * 10^4^, n=8. Values are mean ± s.e.m. Scale bars: 300 μm (ILC, PVN), 375 μm (MPOA), 525 μm (CPu), 600 μm (LS), 750 μm (PIL, A, DMH, PAG) and 30 μm for the insets. **b,** The control injection site included the SN, but not the PIL. In contrast to the injection into the PIL, no fibers were found in the socially implicated brain regions, but there were abundant terminals in the CPu. Scale bars: 300 μm (ILC, PVN), 375 μm (MPOA), 525 μm (CPu), 600 μm (LS), 750 μm (PIL, A, DMH, PAG).

**Figure S3.**
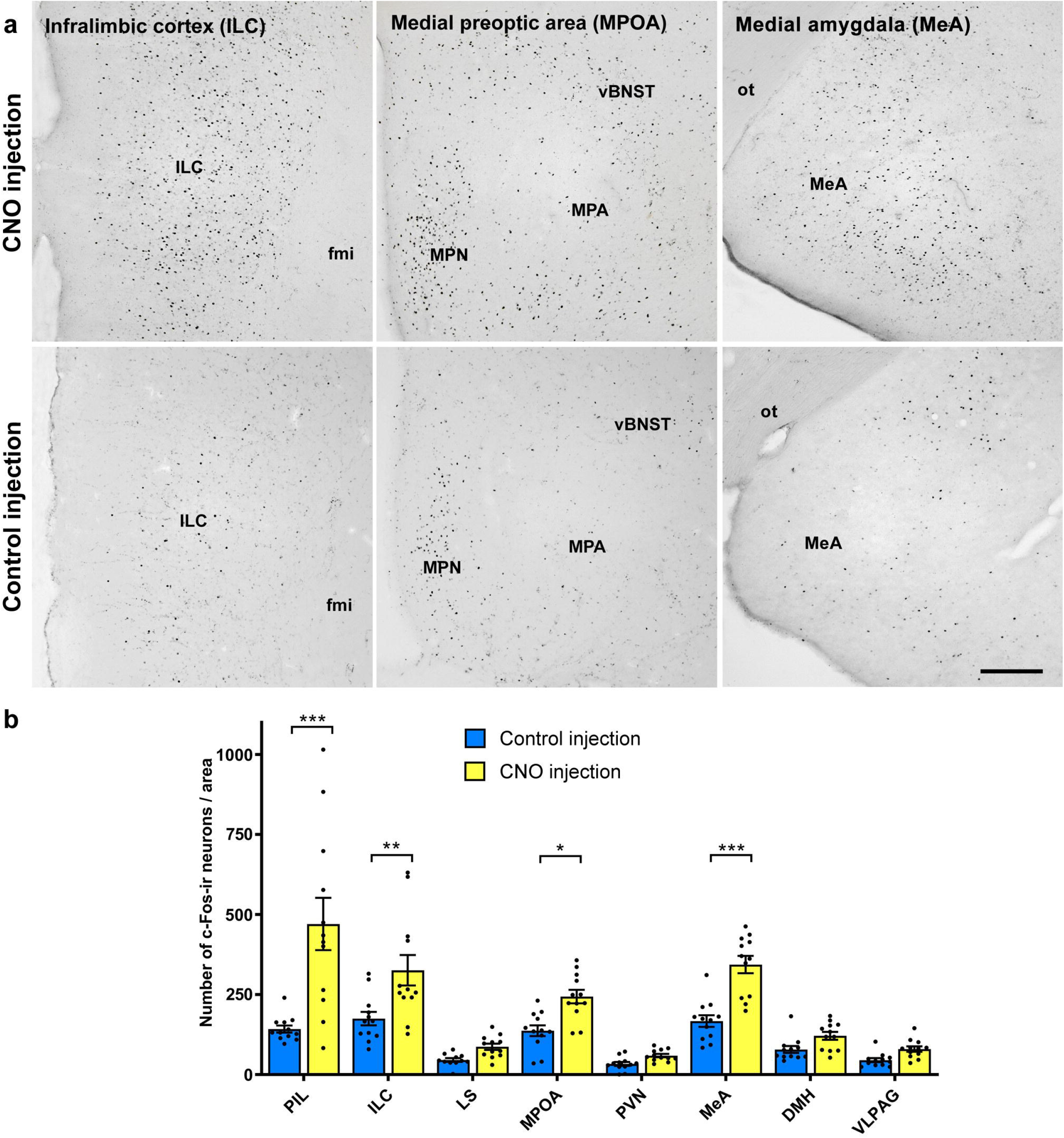
Effect of the chemogenetic stimulation of the PIL on the activity of its target areas. **a,** The upper panels demonstrate the high density of neurons expressing c-Fos at some target areas of the PIL following the chemogenetic stimulation (pAAV-hSyn-hM3D(Gq)-mCherry) of PIL neurons. After control vehicle injection, a markedly lower density of c-Fos activated cells appeared at the same brain regions (bottom panels). Scale bar: 300 μm. **b,** CNO-induced c-Fos activation in the target areas of the PIL. n=12, two-way ANOVA followed by Šídák’s multiple comparisons test, for PIL: *p* < 0.001, for ILC: *p* = 0.0010, for LS: *p* = 0.89, for MPOA: *p* = 0.049, for PVN: *p* = 0.99, for MeA: *p* < 0.001, for DMH: *p* = 0.92, for VLPAG: *p* = 0.97; 0.010 < *\*p* < 0.050, 0.001 < \*\**p* < 0.010 and \*\*\**p* < 0.001. Values are mean ± s.e.m.

**Figure S4.**
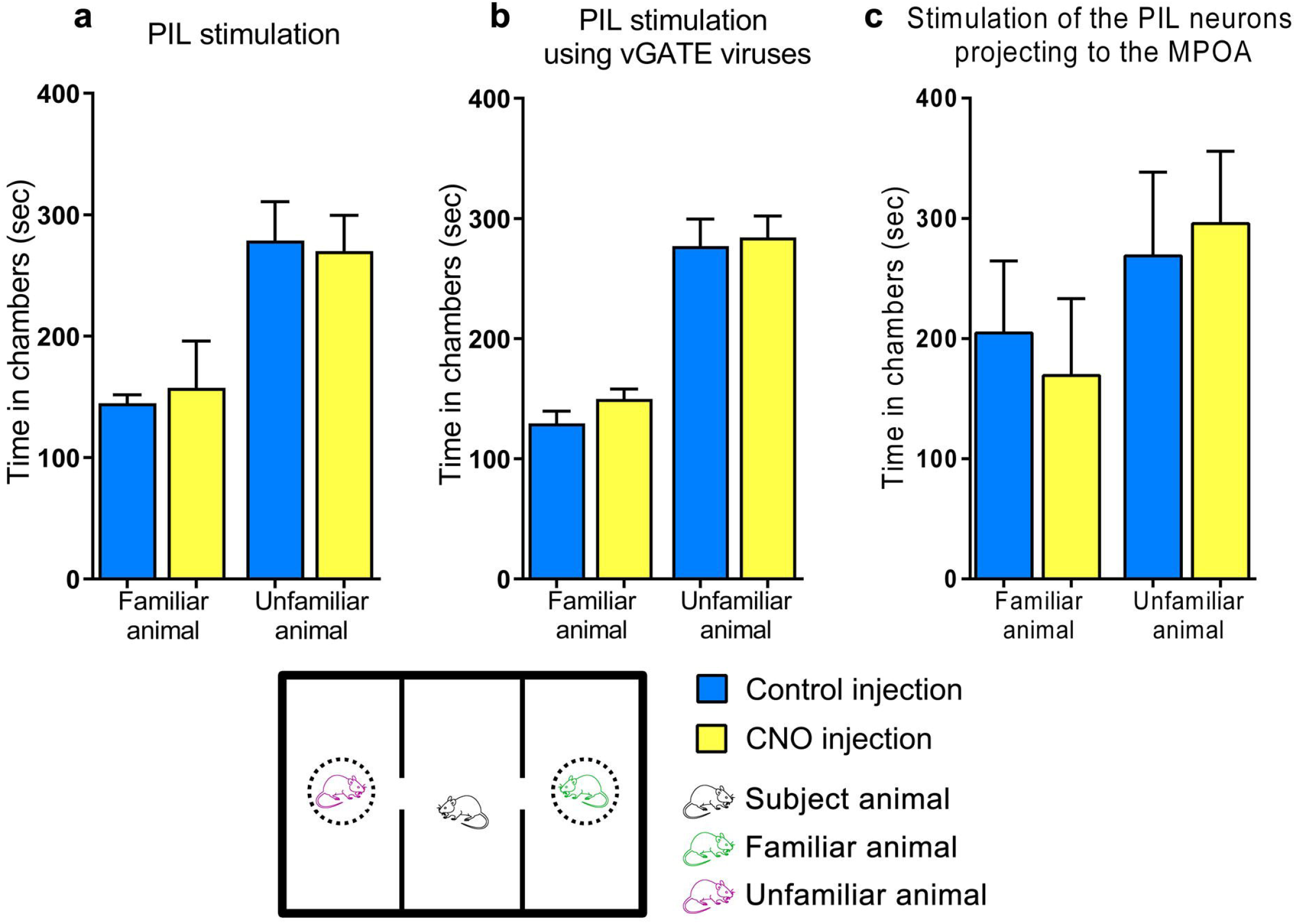
The effect of the chemogenetic stimulation of the PIL on the preference of social novelty. The experiments were carried out in a Three Chambers Sociability Apparatus. The subjects (female rats) could choose between chambers containing a familiar or an unfamiliar female conspecific. **a,** The effect of pAAV-hSyn-hM3D(Gq)-mCherry injected into the PIL. n=5, two-tailed paired t-tests, for familiar animal: *p* = 0.77, for unfamiliar animal: *p* = 0.86. Values are mean ± s.e.m. **b,** vGATE excitatory viral cocktail injected into the PIL: rAAV-(tetO)_7_-**P**_fos_-rtTA; rAAV-**P**_tet_bi-Cre/YC3.60; pAAV-hSyn-DIO-hM3D(Gq)-mCherry. n=12, two-tailed paired t-tests, for familiar animal: *p* = 0.11, for unfamiliar animal: *p* = 0.82. Values are mean ± s.e.m. **c,** Double injections: AAVrg-Ef1a-mCherry-IRES-Cre into the MPOA and pAAV-hSyn-DIO-HA-hM3D(Gq)-IRES-mCitrine into the PIL. n=4, two-tailed paired t-tests, for familiar animal: *p* = 0.70, for unfamiliar animal: *p* = 0.53. Values are mean ± s.e.m.

**Figure S5.**
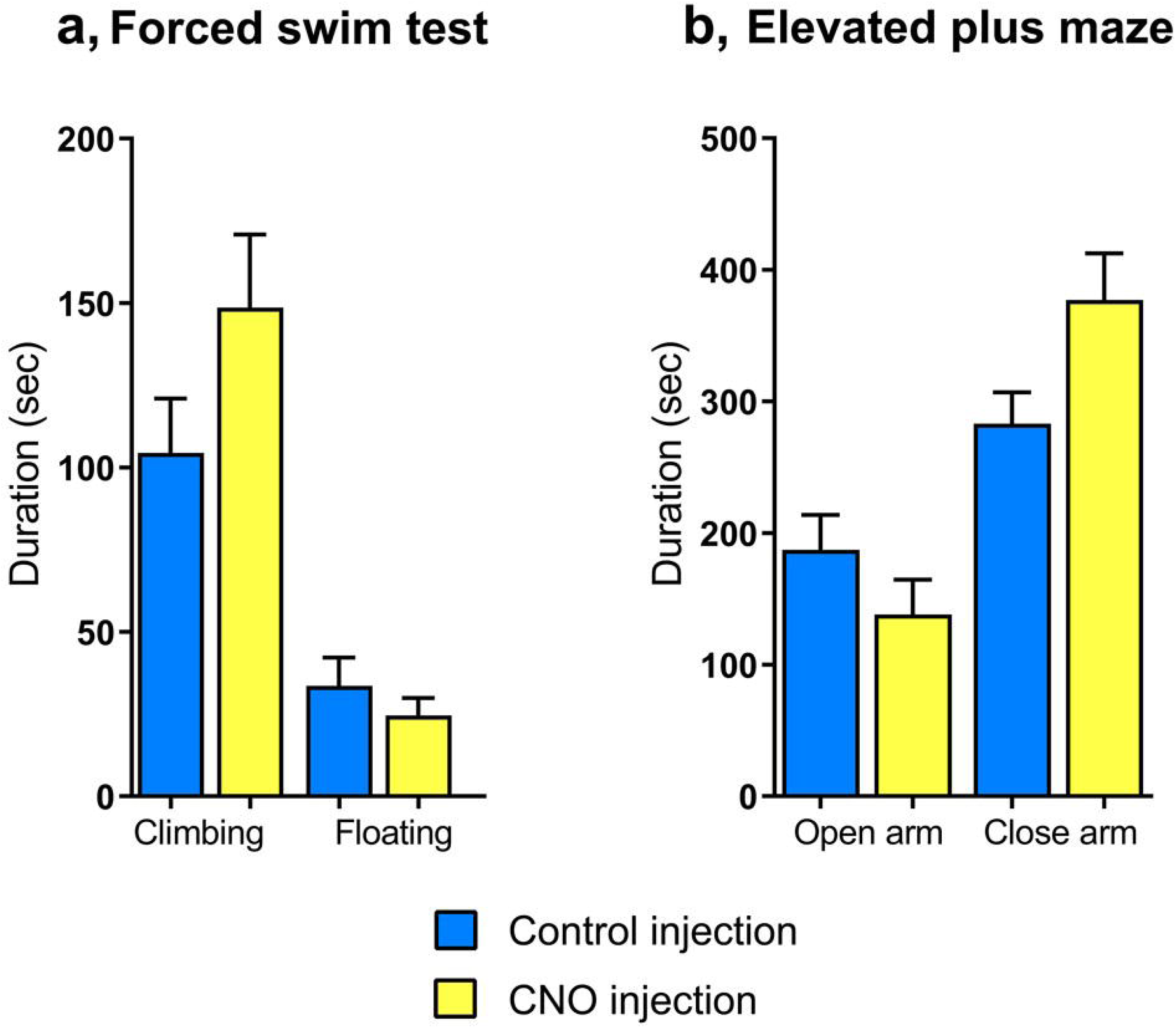
Chemogenetic stimulation of the PIL has no effect on the depression and anxiety-like behaviors of rats. vGATE excitatory viral cocktail injected into the PIL: rAAV-(tetO)_7_-**P**_fos_-rtTA; rAAV-**P**_tet_bi-Cre/YC3.60; pAAV-hSyn-DIO-hM3D(Gq)-mCherry. **a,** In the forced swim test (FST), the time spent with climbing and floating were measure for 6 min. n=6 (each group), two-tailed unpaired t-tests, for climbing: *p* = 0.14, for floating: *p* = 0.40. Values are mean ± s.e.m. **b,** In the elevated plus maze test (EPM), the time spent in the open and close arms were measured for 10 min. n=6 (each group), two-tailed unpaired t-tests, for open arm: *p* = 0.23, for close arm: *p* = 0.073. Values are mean ± s.e.m.

**Figure S6.**
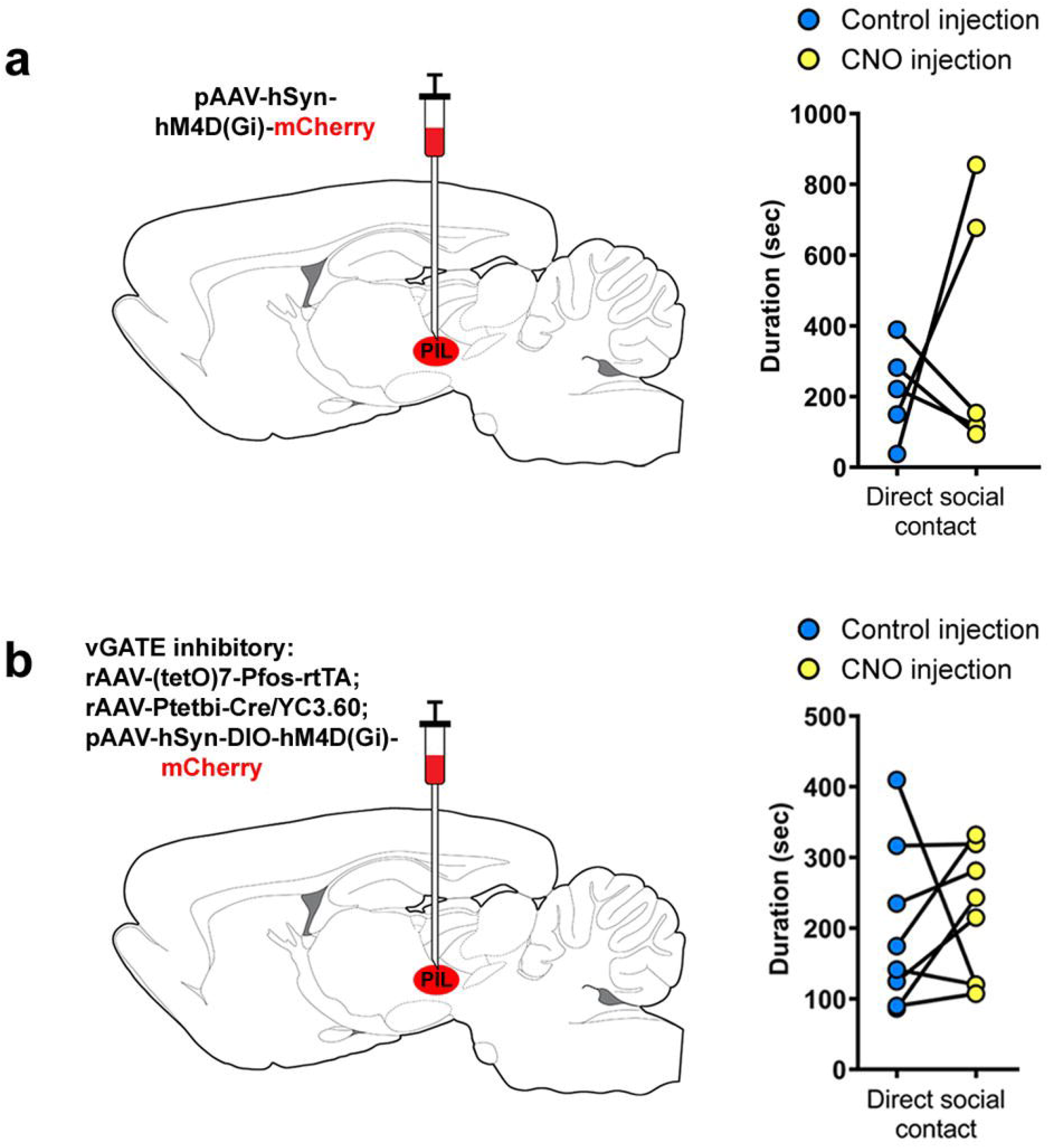
Effect of the chemogenetic inhibition of the PIL on social behavior. The animals were placed in the same cage where they could freely interact with each other. **a,** pAAV-hSyn-hM4D(Gi)-mCherry injected into the PIL. n=5, two-tailed paired t-test, *p* = 0.49. **b,** vGATE inhibitory viral cocktail injected into the PIL: rAAV-(tetO)_7_-**P**_fos_-rtTA; rAAV-**P**_tet_bi-Cre/YC3.60; pAAV-hSyn-DIO-hM4D(Gi)-mCherry. n=8, two-tailed paired t-test, *p* = 0.70.

**Table S1.** Semi-quantitative analysis of labeled fibers in the rat brain following the injection of BDA into the PIL or SN, compared to the distribution of PTH2-containing fibers in the brain. The semi-quantitative analysis of the labeled mCherry-ir fibers represented as none to low (0), moderate (+), high (++), and very high (+++). The last column contains the distribution of PTH2-containing fibers in the brain described previously (Dobolyi *et al*., 2003), which, as we expected, correlates well with the projections of the PIL.

|  | Injection sites |  | PTH2 |
| --- | --- | --- | --- |
|  | SN | PIL |  |
| **Cerebral cortex** |  |  |  |
| Prelimbic cortex | 0 | + | + |
| Infralimbic cortex | 0 | + | + |
| Dorsal peduncular cortex | 0 | + | + |
| Perirhinal cortex | 0 | + | + |
| Frontal cortex | 0 | 0 | 0 |
| Somatomotor cortex | 0 | 0 | 0 |
| Somatosensory cortex (barrel) | 0 | 0 | 0 |
| Cingulate cortex | 0 | 0 | 0 |
| Piriform cortex | 0 | 0 | 0 |
| Entorhinal cortex | 0 | 0 | 0 |
| Ectorhinal cortex | 0 | + | ++ |
| Auditory cortex - primary | 0 | + | + |
| Auditory cortex - secondary | 0 | + | + |
| **Olfactory bulb** |  |  |  |
| Anterior olfactory nucleus | 0 | + | + |
| **Hippocampus** |  |  |  |
| CA1 | 0 | 0 | 0 |
| CA2 | 0 | 0 | 0 |
| CA3 | 0 | 0 | 0 |
| Dentate gyrus | 0 | 0 | 0 |
| Tenia tecta | 0 | 0 | 0 |
| **Septum** |  |  |  |
| Lateral septal nucleus |  |  |  |
| Dorsal | 0 | + | 0 |
| Intermediate | 0 | ++ | ++ |
| Ventral | 0 | +++ | +++ |
| Medial septal nucleus | 0 | + | 0 |
| Subfornical organ | 0 | 0 | 0 |
| **Amygdala** |  |  |  |
| Central nucleus | 0 | ++ | ++ |
| Basolateral nucleus | 0 | + | + |
| Basomedial nucleus | 0 | ++ | + |
| Lateral nucleus |  |  |  |
| Ventral | 0 | 0 | 0 |
| Dorsal | 0 | + | 0 |
| Medial nucleus | 0 | ++ | + |
| Cortical nucleus | 0 | 0 | 0 |
| Amygdalo-striatal transition area | 0 | ++ | ++ |
| **Basal nuclei** |  |  |  |
| Caudate-putamen | +++ | + | + |
| Globus pallidus | ++ | + | 0 |
| Clastrum | 0 | + | 0 |
| Nucleus accumbens |  |  |  |
| Shell | 0 | ++ | + |
| Core | 0 | + | 0 |
| Ventral bed nucleus of the stria terminalis | 0 | +++ | +++ |
| Ventral pallidum | 0 | ++ | ++ |
| **Thalamus** |  |  |  |
| Anterodorsal nucleus | 0 | 0 | 0 |
| Anteroventral nucleus | 0 | 0 | 0 |
| Paraventricular nucleus | 0 | + | + |
| Reticular nucleus | 0 | 0 | 0 |
| Ventral anterior nucleus | 0 | 0 | 0 |
| Ventral lateral nucleus | 0 | 0 | 0 |
| Ventral posterolateral nucleus | 0 | 0 | 0 |
| Ventral posteromedial nucleus | 0 | 0 | 0 |
| Dorsomedial nuclei | 0 | 0 | 0 |
| Posterior nucleus | + | + | + |
| Periventricular grey area | 0 | + | +++ |
| Zona incerta | ++ | +++ | +++ |
| **Hypothalamus** |  |  |  |
| Medial preoptic nucleus | 0 | +++ | +++ |
| Medial preoptic area | 0 | ++ | ++ |
| Lateral preoptic area | 0 | + | + |
| Periventricular nucleus | 0 | ++ | + |
| Supraoptic nucleus | 0 | + | + |
| Suprachiasmatic nucleus | 0 | 0 | 0 |
| Paraventricular nucleus | 0 | ++ | ++ |
| Anterior hypothalamic nucleus | 0 | + | + |
| Arcuate nucleus | 0 | ++ | + |
| Median eminence | 0 | 0 | + |
| Ventromedial nucleus | 0 | ++ | + |
| Dorsomedial nucleus | 0 | +++ | +++ |
| Dorsal hypothalamic area | 0 | ++ | + |
| Posterior hypothalamic nucleus | 0 | ++ | ++ |
| Tuberal part of lateral hypothalamus | 0 | + | + |
| Lateral hypothalamic area | 0 | + | + |
| Supraoptic decussations | 0 | ++ | +++ |
| **Subthalamic nucleus** | + | + | + |
| **Midbrain** |  |  |  |
| Superior colliculus | 0 | + | + |
| Inferior colliculus |  |  |  |
| Central nucleus | 0 | 0 | + |
| External cortex | 0 | + | +++ |
| Periaqueductal central grey |  |  |  |
| Ventrolateral | 0 | ++ | ++ |
| Lateral | 0 | ++ | ++ |
| Dorsolateral | 0 | ++ | ++ |
| Dorsomedial | 0 | ++ | ++ |
| Dorsal raphe nucleus | 0 | + | + |
| Midline raphe nucleus | 0 | 0 | 0 |
| Ventral tegmental area | 0 | 0 | 0 |
| Interpeduncular nucleus | 0 | 0 | 0 |
| Red nucleus | 0 | 0 | 0 |
| Cuneiform nucleus | 0 | +++ | + |
| Lateral parabrachial nucleus | 0 | ++ | ++ |
| Lateral lemniscal nuclei | 0 | ++ | ++ |
| Lateral lemniscus | 0 | + | + |
| Medial paralemniscal nucleus | 0 | + | ++ |
| **Pons** |  |  |  |
| Locus coeruleus | 0 | + | + |
| Dorsal tegmental nucleus | 0 | + | 0 |
| Pontine nuclei | 0 | + | 0 |
| Superior olive | 0 | + | + |
| Pontine reticular formation | + | + | 0 |
| Sensory (principal) trigeminal nucleus | 0 | 0 | 0 |
| Motor trigeminal nucleus | 0 | 0 | 0 |
| Pontine raphe nucleus | 0 | + | + |
| A5 noradrenaline cell group | 0 | 0 | + |
| Vestibular nuclei | 0 | 0 | 0 |
| **Cerebellum** |  |  |  |
| Cortex | 0 | 0 | 0 |
| Nuclei | 0 | 0 | 0 |
| **Medulla oblongata** |  |  |  |
| Cochlear nuclei | 0 | 0 | 0 |
| Gigantocellular reticular nucleus | 0 | + | 0 |
| Parvocellular reticular nucleus | 0 | ++ | + |
| Spinal trigeminal nucleus |  |  |  |
| Interpol part | 0 | + | + |
| Caudal part | 0 | 0 | + |
| Medullary reticular formation | 0 | + | + |
| Inferior olive | 0 | 0 | 0 |
| Medullary raphe nuclei | 0 | 0 | 0 |
| Gracile nucleus | 0 | + | 0 |
| Cuneate nucleus | 0 | + | + |

**Table S2.** Previous anatomical and functional investigations of PIL area. The PIL was named in various ways along the decades: first as the lateral zona incerta or peripeduncular nucleus (PPN), then as parvocellular subparafascicular thalamic nucleus (SPFp) and lately as posterior intralaminar complex of the thalamus. The functions of the area were also investigated, its role in the regulation of maternal behavior, as well as in the auditory system have been determined.

| Name of region | Description | Reference |
| --- | --- | --- |
| Mid-brain area ventromedial to the medial geniculate body | Using microstimulation, the area ventromedial to the medial geniculate body, but not the caudally located lateral midbrain tegmentum, was determined to contain effective injection sites of the milk-ejection reflex. | Tindal and Knaggs (1975) (Tindal and Knaggs, 1975) |
| Lateral zona incerta | Retrograde tracer injected into the supraoptic nucleus labeled in the lateral zona incerta. | Tribollet *et al.* (1985) (Tribollet et al., 1985) |
| Posterior intralaminar nucleus (PIN) | The posterior intralaminar nucleus is a triangular-shaped structure located ventromedial to the medial division of MG and it is retrogradely labeled following injection of axonal markers into the dorsal amygdala. | Ledoux *et al.* (1985) (LeDoux et al., 1985), Ledoux *et al.* (1987) (Ledoux et al., 1987) |
| Peripeduncular nucleus (PPN) | PPN lesions reduced maternal aggression, partially inhibited lactation and caused disturbance in the neuroendocrine control of male and female copulatory behavior. | Hansen and Köhler (1984) (Hansen and Kohler, 1984), López *et al.* (1985) (Lopez and Carrer, 1985a), Factor *et al.* (1993) (Factor et al., 1993) |
| Peripeduncular nucleus (PPN) | Stimuli applied to the PPN evoked potentials in the ventromedial hypothalamic nucleus, which involves synaptic activity in both the lateral amygdaloid nucleus and the bed nucleus of the stria terminalis. | López *et al.* (1985) (Lopez and Carrer, 1985b) |
| Parvicellular division of the subparafascicular and posterior intralaminar nuclei | These auditory-responsive posterior thalamic nuclei project to the medial parvocellular region of the hypothalamic paraventricular nucleus. | Campeau and Watson (2000) (Campeau and Watson, 2000) |
| Posterior intralaminar nucleus (PIN) | As part of the caudal paralaminar thalamic nuclei, PIN plays a role in processing sensory stimuli during emotional situations, to the amygdala. | Linke and Schwegler (2000) (Linke and Schwegler, 2000) |
| Centromedian and parafascicular nuclei (Cm/Pf) | In macaque monkey studies, most Cm/Pf neurons show multimodal sensory activity, responding to auditory, visual and/or somatosensory stimuli. Cm/Pf neurons generally showed greater responses to unexpected sensory stimuli and responded to the sensory stimuli. | Matsumoto *et al.* (2000) (Matsumoto et al., 2001), Saalman (2014) (Saalman, 2014) |
| Intralaminar centromedian/parafascicular thalamic complex (CM/PF) | The CM/PF has been related to motor control and pain, as well as attentional processing, and sexual behavior in animal studies. High-resolution imaging at ultra-high field strength identified activation during attentional processes in the intralaminar CM/PF in humans. | Minamimoto and Kimura (2002) (Minamimoto and Kimura, 2002), Metzger *et al.* |
|  |  | (2013) (Metzger et al., 2013) |
| Subparafascicular and posterior intralaminar nuclei | Subparafascicular and posterior intralaminar nuclei were labeled following the injection of retrograde tracer into the inferior colliculus. | Winer *et al.* (2002) (Winer et al., 2002) |
| Parvocellular subparafascicular thalamic nucleus (SPFp) | The medial subdivision of SPFp contains a dense population of galanin-immunoreactive fibers, originating from galanin neurons in the lumbosacral spinal cord. In contrast, the lateral subdivision contains CGRP-positive fibers and neurons. | Coolen *et al.* (2003) (Coolen et al., 2003) |
| Posterior intralaminar complex of the thalamus (PIL) | A group of PTH2 neurons scattered throughout the posterior intralaminar complex of the thalamus (PIC), an area that includes the parvocellular subparafascicular nucleus, the posterior intralaminar thalamic nucleus, and cells in the most caudal and lateral part of the zona incerta. This area is situated in the rostral end of the lateral mesencephalic tegmentum, demonstrates retrogradely labeled neurons after mediobasal hypothalamic injections and Fos activation in lactating rats. | Palkovits *et al.* (2010)(Palkovits et al., 2010), Cservenák *et al.* (2010)(Cservenak et al., 2010), Cservenák *et al.* (2013)(Cservenak et al., 2013) |
| Posterior intralaminar thalamic nucleus (PIL) | Using expression-based correlation maps and the manual mapping of mouse and human datasets, genes associated with GABAergic neurons (Gad1, Gad2, and Slc32a1), Calbindin 2, Parvalbumin and Insulin-like growth factor binding protein 4 were described in the PIL. | Nagalski *et al.* (2016) (Nagalski et al., 2016) |
| Posterior intralaminar complex of the thalamus (PIL) | The PIL also corresponds to the posterior intralaminar complex, described as containing calbindin but not parvalbumin in the mouse. c-Fos study showed that the PIL is activated upon the social encounter of female rats. The PTH2-positive neurons of PIL innervate and activate the oxytocin cells of PVN. | Cservenák *et al.* (2017)(Cservenak et al., 2017) |
| Posterior intralaminar thalamic nucleus (PIL) | Thalamic cells retrogradely labeled from the lateral amygdala with cholera toxin B subunit mainly located around the auditory thalamus in the PIL and the supragenulate regions. | Barsy *et al.* (2020) (Barsy et al., 2020) |

**Table S3.**
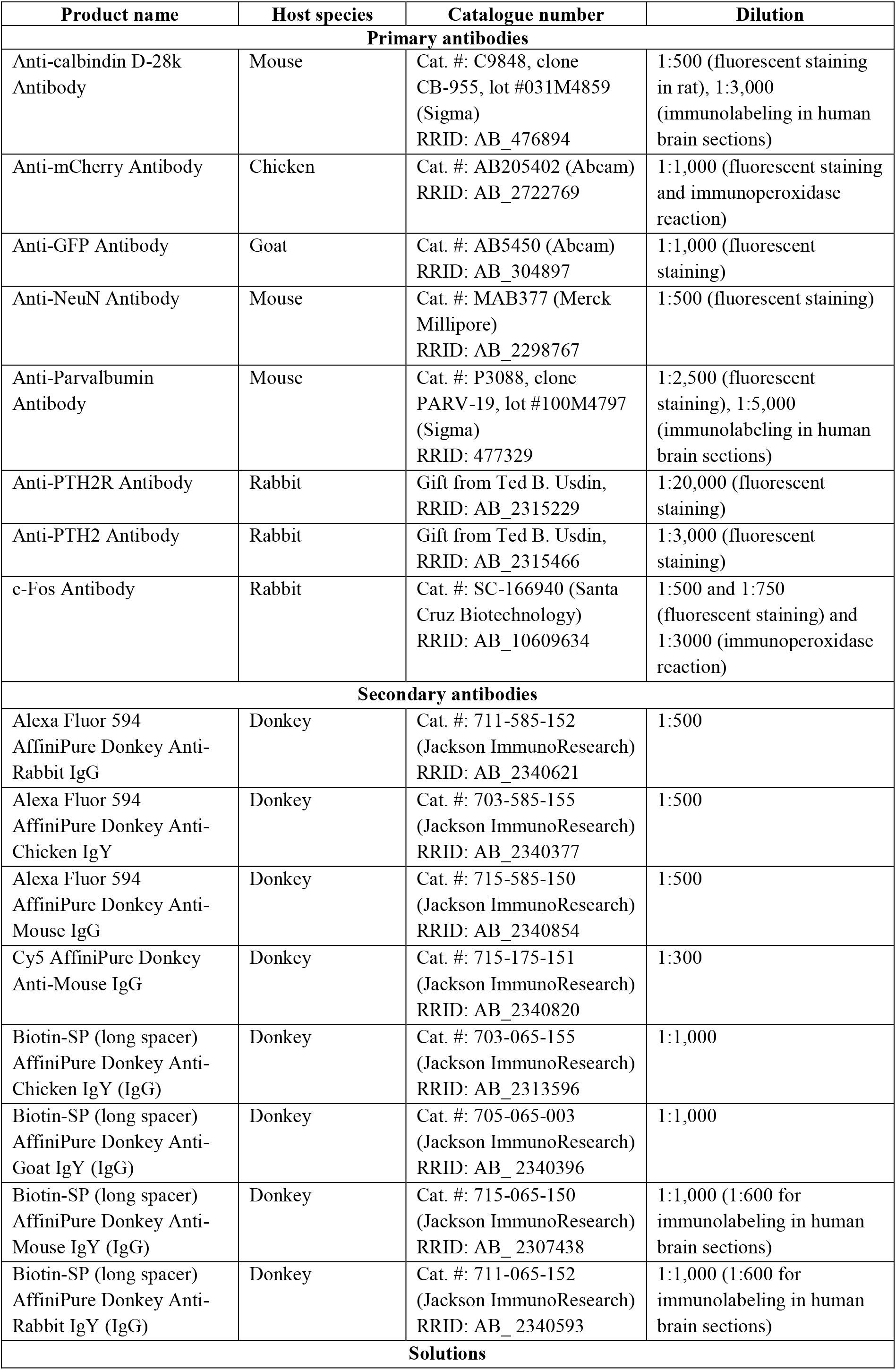
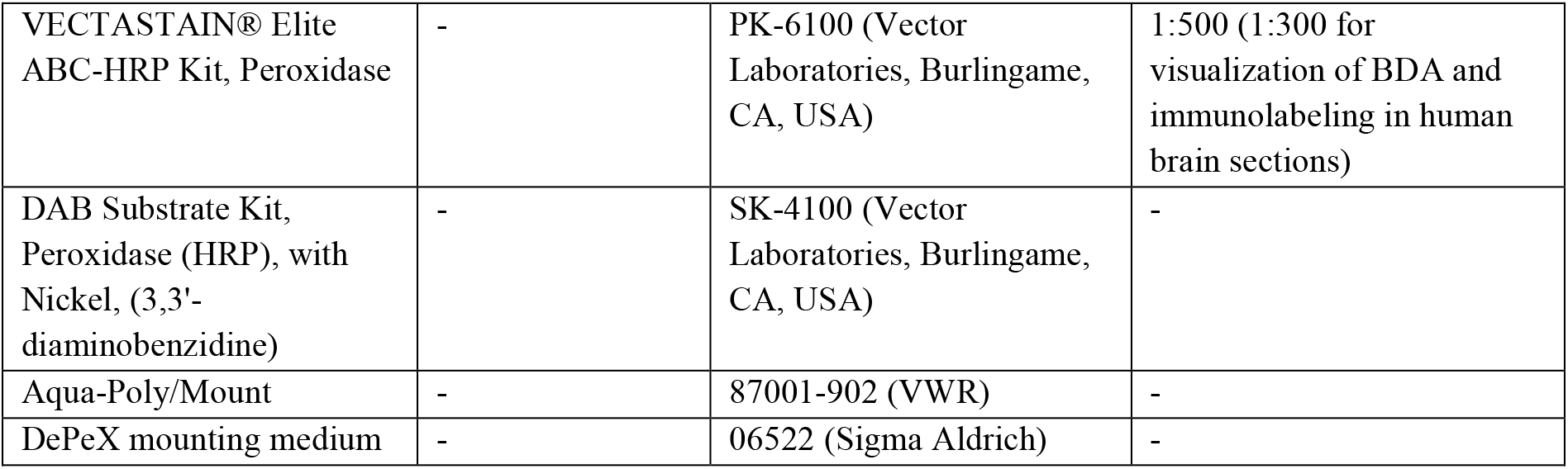
Antibodies and histological reagents used in the study.

| Product name | Host species | Catalogue number | Dilution |
| --- | --- | --- | --- |
| **Primary antibodies** |  |  |  |
| Anti-calbindin D-28k Antibody | Mouse | Cat. #: C9848, clone CB-955, lot #031M4859 (Sigma)<br>RRID: AB_476894 | 1:500 (fluorescent staining in rat), 1:3,000 (immunolabeling in human brain sections) |
| Anti-mCherry Antibody | Chicken | Cat. #: AB205402 (Abcam)<br>RRID: AB_2722769 | 1:1,000 (fluorescent staining and immunoperoxidase reaction) |
| Anti-GFP Antibody | Goat | Cat. #: AB5450 (Abcam)<br>RRID: AB_304897 | 1:1,000 (fluorescent staining) |
| Anti-NeuN Antibody | Mouse | Cat. #: MAB377 (Merck Millipore)<br>RRID: AB_2298767 | 1:500 (fluorescent staining) |
| Anti-Parvalbumin Antibody | Mouse | Cat. #: P3088, clone PARV-19, lot #100M4797 (Sigma)<br>RRID: 477329 | 1:2,500 (fluorescent staining), 1:5,000 (immunolabeling in human brain sections) |
| Anti-PTH2R Antibody | Rabbit | Gift from Ted B. Usdin,<br>RRID: AB_2315229 | 1:20,000 (fluorescent staining) |
| Anti-PTH2 Antibody | Rabbit | Gift from Ted B. Usdin,<br>RRID: AB_2315466 | 1:3,000 (fluorescent staining) |
| c-Fos Antibody | Rabbit | Cat. #: SC-166940 (Santa Cruz Biotechnology)<br>RRID: AB_10609634 | 1:500 and 1:750 (fluorescent staining) and 1:3000 (immunoperoxidase reaction) |
| **Secondary antibodies** |  |  |  |
| Alexa Fluor 594 AffiniPure Donkey Anti-Rabbit IgG | Donkey | Cat. #: 711-585-152 (Jackson ImmunoResearch)<br>RRID: AB_2340621 | 1:500 |
| Alexa Fluor 594 AffiniPure Donkey Anti-Chicken IgY | Donkey | Cat. #: 703-585-155 (Jackson ImmunoResearch)<br>RRID: AB_2340377 | 1:500 |
| Alexa Fluor 594 AffiniPure Donkey Anti-Mouse IgG | Donkey | Cat. #: 715-585-150 (Jackson ImmunoResearch)<br>RRID: AB_2340854 | 1:500 |
| Cy5 AffiniPure Donkey Anti-Mouse IgG | Donkey | Cat. #: 715-175-151 (Jackson ImmunoResearch)<br>RRID: AB_2340820 | 1:300 |
| Biotin-SP (long spacer) AffiniPure Donkey Anti-Chicken IgY (IgG) | Donkey | Cat. #: 703-065-155 (Jackson ImmunoResearch)<br>RRID: AB_2313596 | 1:1,000 |
| Biotin-SP (long spacer) AffiniPure Donkey Anti-Goat IgY (IgG) | Donkey | Cat. #: 705-065-003 (Jackson ImmunoResearch)<br>RRID: AB_2340396 | 1:1,000 |
| Biotin-SP (long spacer) AffiniPure Donkey Anti-Mouse IgY (IgG) | Donkey | Cat. #: 715-065-150 (Jackson ImmunoResearch)<br>RRID: AB_2307438 | 1:1,000 (1:600 for immunolabeling in human brain sections) |
| Biotin-SP (long spacer) AffiniPure Donkey Anti-Rabbit IgY (IgG) | Donkey | Cat. #: 711-065-152 (Jackson ImmunoResearch)<br>RRID: AB_2340593 | 1:1,000 (1:600 for immunolabeling in human brain sections) |
| **Solutions** |  |  |  |
| VECTASTAIN® Elite ABC-HRP Kit, Peroxidase | - | PK-6100 (Vector Laboratories, Burlingame, CA, USA) | 1:500 (1:300 for visualization of BDA and immunolabeling in human brain sections) |
| DAB Substrate Kit, Peroxidase (HRP), with Nickel, (3,3'-diaminobenzidine) | - | SK-4100 (Vector Laboratories, Burlingame, CA, USA) | - |
| Aqua-Poly/Mount | - | 87001-902 (VWR) | - |
| DePeX mounting medium | - | 06522 (Sigma Aldrich) | - |

**Table S4.**
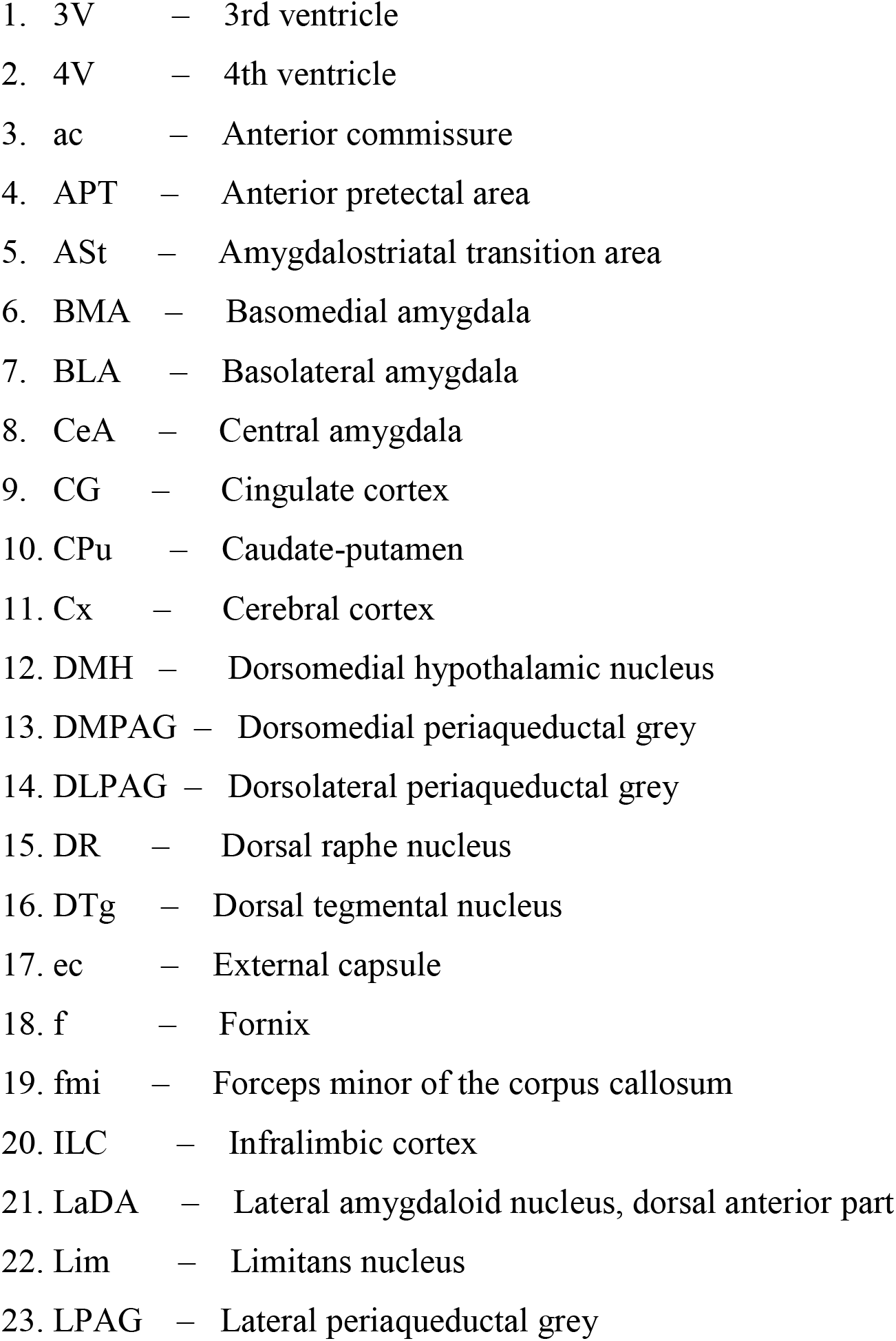
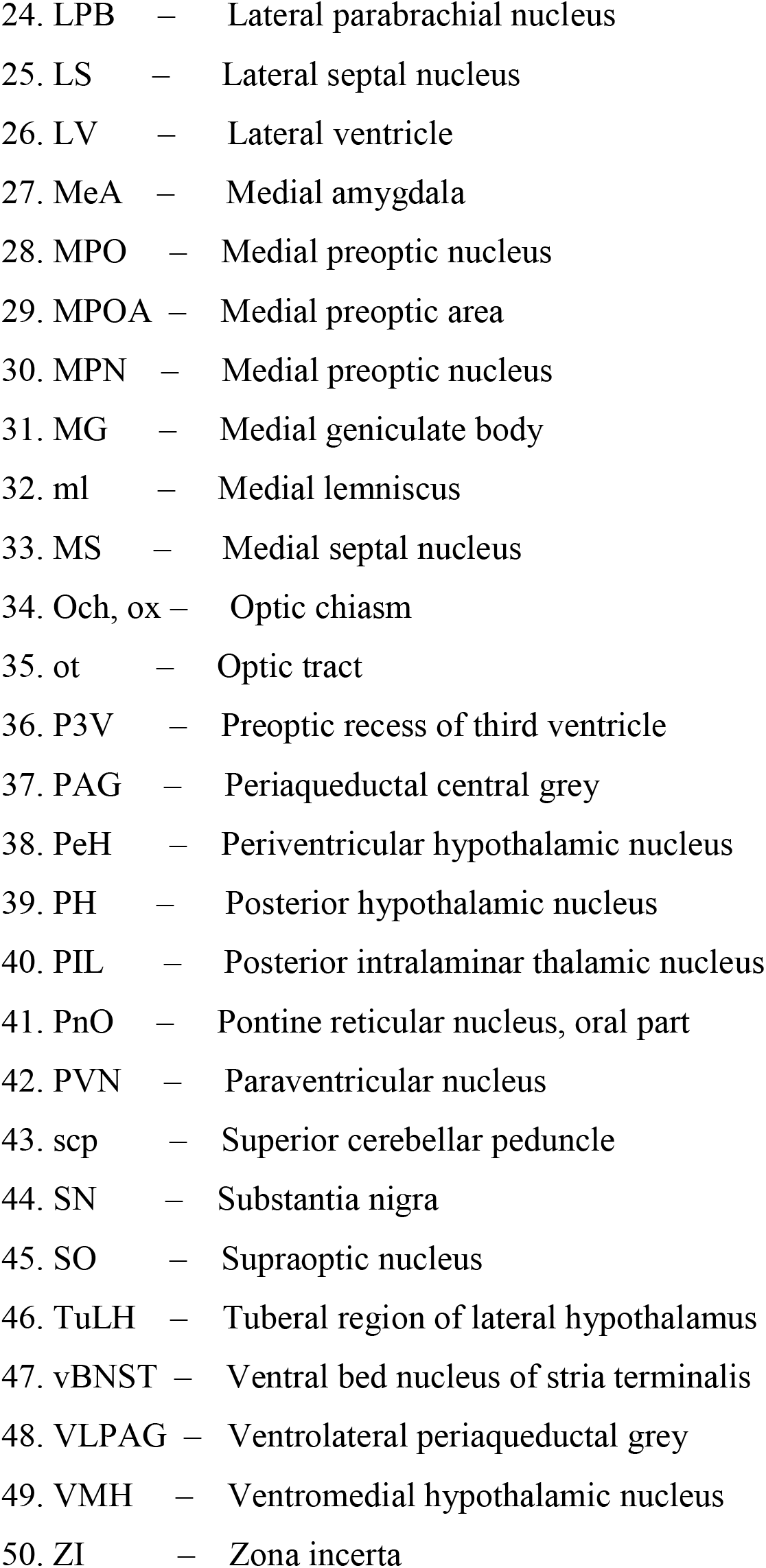
List of abbreviations.

## References

Ahmadlou, M., Houba, J.H.W., van Vierbergen, J.F.M., Giannouli, M., Gimenez, G.A., van Weeghel, C., Darbanfouladi, M., Shirazi, M.Y., Dziubek, J., Kacem, M., de Winter, F., and Heimel, J.A. (2021). A cell type-specific cortico-subcortical brain circuit for investigatory and novelty-seeking behavior. Science 372.

Anneser, L., Alcantara, I.C., Gemmer, A., Mirkes, K., Ryu, S., and Schuman, E.M. (2020). The neuropeptide Pth2 dynamically senses others via mechanosensation. Nature 588, 653–657.

Bankhead, P., Loughrey, M.B., Fernandez, J.A., Dombrowski, Y., McArt, D.G., Dunne, P.D., McQuaid, S., Gray, R.T., Murray, L.J., Coleman, H.G., James, J.A., Salto-Tellez, M., and Hamilton, P.W. (2017). QuPath: Open source software for digital pathology image analysis. Sci Rep 7, 16878.

Barsy, B., Kocsis, K., Magyar, A., Babiczky, A., Szabo, M., Veres, J.M., Hillier, D., Ulbert, I., Yizhar, O., and Matyas, F. (2020). Associative and plastic thalamic signaling to the lateral amygdala controls fear behavior. Nat Neurosci 23, 625–637.

Campeau, S., and Watson, S.J., Jr. (2000). Connections of some auditory-responsive posterior thalamic nuclei putatively involved in activation of the hypothalamo-pituitary-adrenocortical axis in response to audiogenic stress in rats: an anterograde and retrograde tract tracing study combined with Fos expression. J Comp Neurol 423, 474–491.

Coolen, L.M., Veening, J.G., Petersen, D.W., and Shipley, M.T. (2003). Parvocellular subparafascicular thalamic nucleus in the rat: anatomical and functional compartmentalization. J Comp Neurol 463, 117–131.

Cservenak, M., Bodnar, I., Usdin, T.B., Palkovits, M., Nagy, G.M., and Dobolyi, A. (2010). Tuberoinfundibular peptide of 39 residues is activated during lactation and participates in the suckling-induced prolactin release in rat. Endocrinology 151, 5830–5840.

Cservenak, M., Keller, D., Kis, V., Fazekas, E.A., Ollos, H., Leko, A.H., Szabo, E.R., Renner, E., Usdin, T.B., Palkovits, M., and Dobolyi, A. (2017). A thalamo-hypothalamic pathway that activates oxytocin neurons in social contexts in female rats. Endocrinology 158, 335–348.

Cservenak, M., Szabo, E.R., Bodnar, I., Leko, A., Palkovits, M., Nagy, G.M., Usdin, T.B., and Dobolyi, A. (2013). Thalamic neuropeptide mediating the effects of nursing on lactation and maternal motivation. Psychoneuroendocrinology 38, 3070–3084.

Deschrijver, E., Wiersema, J.R., and Brass, M. (2017). Action-based touch observation in adults with high functioning autism: Can compromised self-other distinction abilities link social and sensory everyday problems? Soc Cogn Affect Neurosci 12, 273–282.

Dimen, D., Puska, G., Szendi, V., Sipos, E., Zelena, D., and Dobolyi, A. (2021). Sex-specific parenting and depression evoked by preoptic inhibitory neurons. iScience 24, 103090.

Dobolyi, A. (2009). Central amylin expression and its induction in rat dams. J Neurochem 111, 1490–1500.

Dobolyi, A., Cservenak, M., and Young, L.J. (2018). Thalamic integration of social stimuli regulating parental behavior and the oxytocin system. Front Neuroendocrinol 51, 102–115.

Dobolyi, A., Palkovits, M., and Usdin, T.B. (2010). The TIP39-PTH2 receptor system: unique peptidergic cell groups in the brainstem and their interactions with central regulatory mechanisms. Prog Neurobiol 90, 29–59.

Dobolyi, A., Ueda, H., Uchida, H., Palkovits, M., and Usdin, T.B. (2002). Anatomical and physiological evidence for involvement of tuberoinfundibular peptide of 39 residues in nociception. Proc Natl Acad Sci U S A 99, 1651–1656.

Ebbesen, C.L., and Froemke, R.C. (2021). Body language signals for rodent social communication. Curr Opin Neurobiol 68, 91–106.

Ebert, D.H., and Greenberg, M.E. (2013). Activity-dependent neuronal signalling and autism spectrum disorder. Nature 493, 327–337.

Ellingsen, D.M., Leknes, S., Loseth, G., Wessberg, J., and Olausson, H. (2015). The Neurobiology Shaping Affective Touch: Expectation, Motivation, and Meaning in the Multisensory Context. Front Psychol 6, 1986.

Factor, E.M., Mayer, A.D., and Rosenblatt, J.S. (1993). Peripeduncular nucleus lesions in the rat: I. Effects on maternal aggression, lactation, and maternal behavior during pre- and postpartum periods. Behav Neurosci 107, 166–185.

Farkas, I., Kallo, I., Deli, L., Vida, B., Hrabovszky, E., Fekete, C., Moenter, S.M., Watanabe, M., and Liposits, Z. (2010). Retrograde endocannabinoid signaling reduces GABAergic synaptic transmission to gonadotropin-releasing hormone neurons. Endocrinology 151, 5818–5829.

Fenno, L.E., Mattis, J., Ramakrishnan, C., Hyun, M., Lee, S.Y., He, M., Tucciarone, J., Selimbeyoglu, A., Berndt, A., Grosenick, L., Zalocusky, K.A., Bernstein, H., Swanson, H., Perry, C., Diester, I., Boyce, F.M., Bass, C.E., Neve, R., Huang, Z.J., and Deisseroth, K. (2014). Targeting cells with single vectors using multiple-feature Boolean logic. Nat Methods 11, 763–772.

Fernandes, A.M., Beddows, E., Filippi, A., and Driever, W. (2013). Orthopedia transcription factor otpa and otpb paralogous genes function during dopaminergic and neuroendocrine cell specification in larval zebrafish. PLoS One 8, e75002.

Hansen, S., and Kohler, C. (1984). The importance of the peripeduncular nucleus in the neuroendocrine control of sexual behavior and milk ejection in the rat. Neuroendocrinology 39, 563–572.

Hasan, M.T., Althammer, F., Silva da Gouveia, M., Goyon, S., Eliava, M., Lefevre, A., Kerspern, D., Schimmer, J., Raftogianni, A., Wahis, J., Knobloch-Bollmann, H.S., Tang, Y., Liu, X., Jain, A., Chavant, V., Goumon, Y., Weislogel, J.M., Hurlemann, R., Herpertz, S.C., Pitzer, C., Darbon, P., Dogbevia, G.K., Bertocchi, I., Larkum, M.E., Sprengel, R., Bading, H., Charlet, A., and Grinevich, V. (2019). A Fear Memory Engram and Its Plasticity in the Hypothalamic Oxytocin System. Neuron 103, 133–146.

Kuo, J., and Usdin, T.B. (2007). Development of a rat parathyroid hormone 2 receptor antagonist. Peptides 28, 887–892.

Ledoux, J.E., Ruggiero, D.A., Forest, R., Stornetta, R., and Reis, D.J. (1987). Topographic organization of convergent projections to the thalamus from the inferior colliculus and spinal cord in the rat. J Comp Neurol 264, 123–146.

LeDoux, J.E., Ruggiero, D.A., and Reis, D.J. (1985). Projections to the subcortical forebrain from anatomically defined regions of the medial geniculate body in the rat. J Comp Neurol 242, 182–213.

Linke, R., and Schwegler, H. (2000). Convergent and complementary projections of the caudal paralaminar thalamic nuclei to rat temporal and insular cortex. Cereb Cortex 10, 753–771.

Lopez, H.S., and Carrer, H.F. (1985a). Evidence for peripeduncular neurons having a role in the control of feminine sexual behavior in the rat. Physiol Behav 35, 205–208.

Lopez, H.S., and Carrer, H.F. (1985b). Further studies on peripeduncular-hypothalamic pathways involved in sexual behavior in the female rat. Exp Neurol 88, 241–252.

Mai, J.K. (2008). Atlas of the Human Brain (Academic Press).

Matsumoto, N., Minamimoto, T., Graybiel, A.M., and Kimura, M. (2001). Neurons in the thalamic CM-Pf complex supply striatal neurons with information about behaviorally significant sensory events. J Neurophysiol 85, 960–976.

McIlmoyl, M. (1965). Two neurological staining techniques utilizing the dye luxol fast blue. Can J Med Technol 27, 118–123.

Metzger, C.D., van der Werf, Y.D., and Walter, M. (2013). Functional mapping of thalamic nuclei and their integration into cortico-striatal-thalamo-cortical loops via ultra-high resolution imaging-from animal anatomy to in vivo imaging in humans. Front Neurosci 7, 24.

Minamimoto, T., and Kimura, M. (2002). Participation of the thalamic CM-Pf complex in attentional orienting. J Neurophysiol 87, 3090–3101.

Nagalski, A., Puelles, L., Dabrowski, M., Wegierski, T., Kuznicki, J., and Wisniewska, M.B. (2016). Molecular anatomy of the thalamic complex and the underlying transcription factors. Brain Struct Funct 221, 2493–2510.

Palkovits, M., Usdin, T.B., Makara, G.B., and Dobolyi, A. (2010). Tuberoinfundibular peptide of 39 residues-immunoreactive fibers in the zona incerta and the supraoptic decussations terminate in the neuroendocrine hypothalamus. Neurochem Res 35, 2078–2085.

Paxinos, G., and Watson, C. (2007). The rat brain in stereotaxic coordinates (San Diego: Academic Press).

Saalmann, Y.B. (2014). Intralaminar and medial thalamic influence on cortical synchrony, information transmission and cognition. Front Syst Neurosci 8, 83.

Shamay-Tsoory, S.G., and Eisenberger, N.I. (2021). Getting in touch: A neural model of comforting touch. Neurosci Biobehav Rev 130, 263–273.

Tang, Y., Benusiglio, D., Lefevre, A., Hilfiger, L., Althammer, F., Bludau, A., Hagiwara, D., Baudon, A., Darbon, P., Schimmer, J., Kirchner, M.K., Roy, R.K., Wang, S., Eliava, M., Wagner, S., Oberhuber, M., Conzelmann, K.K., Schwarz, M., Stern, J.E., Leng, G., Neumann, I.D., Charlet, A., and Grinevich, V. (2020). Social touch promotes interfemale communication via activation of parvocellular oxytocin neurons. Nat Neurosci 23, 1125–1137.

Tindal, J.S., and Knaggs, G.S. (1975). Further studies on the afferent path of the milk-ejection reflex in the brain stem of the rabbit. J Endocrinol 66, 107–113.

Tribollet, E., Armstrong, W.E., Dubois-Dauphin, M., and Dreifuss, J.J. (1985). Extra-hypothalamic afferent inputs to the supraoptic nucleus area of the rat as determined by retrograde and anterograde tracing techniques. Neuroscience 15, 135–148.

Tsuneoka, Y., and Funato, H. (2021). Cellular Composition of the Preoptic Area Regulating Sleep, Parental, and Sexual Behavior. Front Neurosci 15, 649159.

Usdin, T.B., Hoare, S.R., Wang, T., Mezey, E., and Kowalak, J.A. (1999). TIP39: a new neuropeptide and PTH2-receptor agonist from hypothalamus. Nat Neurosci 2, 941–943.

Wang, J., Palkovits, M., Usdin, T.B., and Dobolyi, A. (2006). Forebrain projections of tuberoinfundibular peptide of 39 residues (TIP39)-containing subparafascicular neurons. Neuroscience 138, 1245–1263.

Wang, T., Palkovits, M., Rusnak, M., Mezey, E., and Usdin, T.B. (2000). Distribution of parathyroid hormone-2 receptor-like immunoreactivity and messenger RNA in the rat nervous system. Neuroscience 100, 629–649.

Winer, J.A., Chernock, M.L., Larue, D.T., and Cheung, S.W. (2002). Descending projections to the inferior colliculus from the posterior thalamus and the auditory cortex in rat, cat, and monkey. Hear Res 168, 181–195.

Zelikowsky, M., Hui, M., Karigo, T., Choe, A., Yang, B., Blanco, M.R., Beadle, K., Gradinaru, V., Deverman, B.E., and Anderson, D.J. (2018). The Neuropeptide Tac2 Controls a Distributed Brain State Induced by Chronic Social Isolation Stress. Cell 173, 1265–1279.

Zilkha, N., Sofer, Y., Kashash, Y., and Kimchi, T. (2021). The social network: Neural control of sex differences in reproductive behaviors, motivation, and response to social isolation. Curr Opin Neurobiol 68, 137–151.

